# Pan-primate DNA methylation clocks

**DOI:** 10.1101/2020.11.29.402891

**Authors:** Steve Horvath, Amin Haghani, Joseph A. Zoller, Ake T. Lu, Jason Ernst, Matteo Pellegrini, Anna J. Jasinska, Julie A. Mattison, Adam B. Salmon, Ken Raj, Markus Horvath, Kimberly C. Paul, Beate R. Ritz, Todd R. Robeck, Maria Spriggs, Erin E. Ehmke, Susan Jenkins, Cun Li, Peter W. Nathanielsz

## Abstract

DNA methylation data have been successfully used to develop highly accurate estimators of age (“epigenetic clocks”) in many mammalian species. With a view of extending such epigenetic clocks to all primate species, we analyzed DNA methylation profiles of 2400 tissues derived from 37 primate species including 11 haplorhine species (baboons, marmosets, vervets, rhesus macaque, chimpanzees, gorillas, orangutan, humans) and 26 strepsirrhine species (suborders Lemuriformes and Lorisiformes). From these we present here, pan-primate epigenetic clocks which are highly accurate for all primates including humans (age correlation R=0.98). We also carried out in-depth analysis of baboon DNA methylation profiles and generated five epigenetic clocks for baboons (Olive-yellow baboon hybrid), one of which, the pan-tissue epigenetic clock, was trained on seven tissue types (fetal cerebral cortex, adult cerebral cortex, cerebellum, adipose, heart, liver, and skeletal muscle) with ages ranging from late fetal life to 22.8 years of age. To facilitate translation of findings in baboons to humans, we further constructed two dual-species, human-baboon clocks. We also identified and present here, epigenetic predictors of sex that apply to all primate species. Low overlap can be observed between age- and sex-related CpGs. Overall, this study advances our understanding of conserved age- and sex-related epigenetic changes in primates, and provides biomarkers to study the aging of all primate species with the facility to readily translate any findings between primate species.

## INTRODUCTION

During development, germline DNA methylation is erased, and then re-established in tissue-specific patterns as the developmental program unfolds after implantation [1]. The primary role of DNA methylation is to regulate the expression of genes, ensuring the production of appropriate ones in different tissues and organs. Our understanding of age related changes in cytosine methylation has progressed rapidly [2-4] because of the technical advancement of methylation array platforms which simultaneously quantify thousands of individual CpGs at known locations on the human genome. It was observed that age-based methylation changes accompany the functional decline of adult stem cells and that even small methylation changes can lead to loss of regulatory control of transcription, either directly or via additive effects [5, 6].

Crucially, the correlation between methylation changes of some CpGs with chronological age over the course of an entire lifespan was found to be very strong [7, 8]. This allowed methylation levels of several CpGs to be combined and developed into an accurate age estimators, which are reviewed in [8, 9]. An example of this is the human pan-tissue epigenetic age estimator which combines the weighted methylation average of 353 CpGs, into an age estimate that is referred to as DNAm age or epigenetic age [10]. The discrepancy between epigenetic age and chronological age, termed epigenetic age acceleration, is associated with mortality risk and a wide range of age-related conditions [11, 12]. This implies that epigenetic clocks successfully capture biological age, if not in its entirety, then at least to some measurable level.

First generation epigenetic clocks measured chronological age while second generation clocks were designed to assess mortality/morbidity risk [8, 13, 14]. Here we present third generation epigenetic clocks that measure age in multiple species based on a single formula (regression model). Toward this end, we measure cytosine methylation levels in 37 primate species including strepsirrhines and haplorhines that diverged shortly after the emergence of the first true primates.

For ethical and practical reasons, anti-aging studies are difficult in great apes. The olive-yellow hybrid baboons living in the Southwest National Primate Research Center (SNPRC) can be employed to study biological changes including age-related processes [15]. Although baboons are an attractive primate model for studying aging, their lifespan, while considerably shorter than that of humans (max. 122.5 years), is still substantial (max. 37.5 years) *[16]*, and leads to high maintenance costs, especially for studies on anti-aging interventions that use longevity as the primary measure of effect. The presented baboon clocks promise to greatly shorten the duration of baboon studies and hence lower the burden and costs associated with evaluating anti-aging interventions in this important primate species.

In previous publications, we have presented epigenetic clocks for the vervet monkey [17], rhesus macaque [18], and common marmosets [19]. Here we present epigenetic clocks for baboons, lemurs, and other considered primate species.

To test whether sex modulates epigenetic aging effects, we determine the overlap between age- and sex-associated CpGs in different primate species and tissues.

This article addressed four goals. First, to develop epigenetic clocks for baboons (olive-yellow baboon hybrids) and dual species human-baboon clocks. Second, to develop epigenetic clocks that apply to all primate species. Toward this end we leverage data from both strepsirrhines (e.g. lemurs) and haplorhines (e.g. humans) which diverged about 74 million years ago according to molecular estimates based on mitochondrial genomes [20, 21]. Third, to evaluate the effect of age on individual cytosine methylation levels in different tissues and primate species. Fourth, to study the relationship between sex and age on cytosine methylation across primate species.

## Results

We generated DNA methylation data from n=2398 tissue samples from 37 different primate species (**Supplementary Table S1**) including 26 strepsirrhine primate species (suborders Lemuriformes and Lorisiformes). To facilitate cross species comparisons, we used the mammalian methylation array that applies to all mammalian species [22]. The n=326 samples from baboon tissues (fetal cerebral cortex, adult heart, adipose, cerebellum, cerebral cortex, liver, and skeletal muscle) come from baboons with comparable age distributions (**Supplementary Table S2**), which facilitates the comparison of age effects across tissues. Unsupervised hierarchical clustering of the baboon samples reveals that the samples cluster largely by tissue type (**Supplementary Figure S1**).

### Epigenetic clocks for baboons

To arrive at unbiased estimates of the epigenetic clocks, we carried out cross-validation analyses of the training datasets. This generates unbiased estimates of the age correlation R (defined as Pearson correlation between the estimated age (DNAm age) and chronological age) as well as the median absolute error, which indicates concordance of the estimated age with chronological age. The different baboon epigenetic clocks that we constructed differed from each other with regards to tissue type, species, and measure of age. Some clocks apply to all tissues (pan-tissue clocks, **Figure 1A**) while others are tailor-made for specific tissues/organs and named accordingly. We developed two brain-specific epigenetic clocks for baboons: one was trained using two brain regions (cerebellum and cerebral cortex, R=0.96, **Figure 1B, Supplementary Figure S2**) and the other was only trained on cerebral cortex samples (R=0.97, **Figure 1C**). The baboon pan-tissue clock, which was trained on all available tissues, is highly accurate in age estimation of all the different baboon tissue samples (R=0.96 and median error 1.1 years, **Figure 1A, Supplementary Figure S3**), including cerebellum (R=0.93, **Supplementary Figure S3C**) and cerebral cortex samples (R=0.99, **Supplementary Figure S3D**). Although all of these different clocks exhibit high age correlations, pan tissue clocks may have different biological properties from tissue specific clocks, e.g. pan-tissue clocks tend to be less correlated with changes in cellular composition [8].

**Figure 1:**
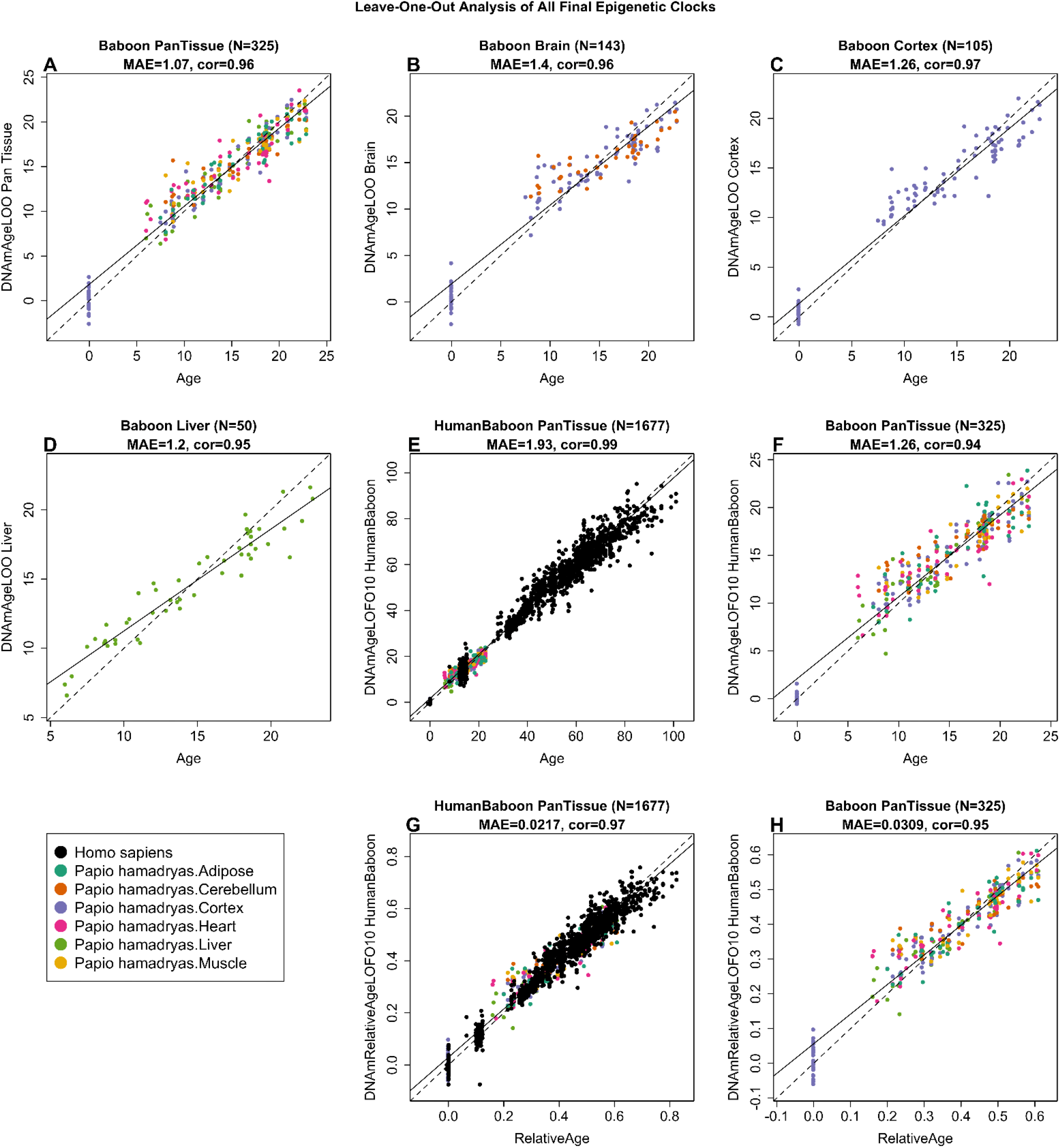
Cross-validation study of epigenetic clocks for baboons and humans. A-C) Three epigenetic clocks that were trained on baboon tissues: A) all tissues, B) all brain tissues, C) cerebral. D,E) Human-baboon clock for relative age applied to D) both species and E) baboons only. Relative age was defined as the ratio of chronological age to maximum lifespan. D) Human samples are colored in (black) and baboon samples (colored by tissue type). Each panel reports the sample size, correlation coefficient, median absolute error (MAE). Dots are colored by tissue type or species. A-C) “LOO” denotes the leave-one-out cross validation estimates of DNA methylation age (y-axis, in units of years). D,E) “LOFO10” denotes the ten-fold cross-validation estimates of age (y-axis, in years).

To generate hybrid human-baboon clocks, human DNA methylation profiles were added to the baboon DNA methylation profiles in the training dataset. From these, two human-baboon pan-tissue clocks were developed. The human-baboon clock for *age* estimates age estimates age in units of years (**Figure 1E,F**). The cross validation analysis indicates that the human-baboon clock for age is highly correlated with age across both tissues from both species (R=0.99, **Figure 1E**) and across baboon tissues (R=0.94, **Figure 1F**).

The human-baboon clock for *relative age* estimates the ratio of chronological age to maximum lifespan of the species, with values between 0 and 1. This ratio allows alignment and biologically meaningful comparison between species with different average and maximal lifespans (baboon, 37.5 years and human, 122.5 years), which is not afforded by mere measurement of absolute age. The human-baboon clock for relative age is highly accurate when applied to both species together (R=0.97, **Figure 1G**) and only marginally less so when the analysis is restricted to baboon tissues (R=0.95, **Figure 1H**).

To determine the cross-tissue performance and the cross-species applicability of the baboon clocks, we applied them to an array of tissues from key human organs (n=1352 from 16 tissues). The application of pan-tissue baboon clock to *human* tissues produced epigenetic age estimates that are poorly calibrated; meaning that they exhibited poor concordance with chronological age (high median absolute error), but they nevertheless showed low to moderate correlations with ages with several human tissues (e.g. human adipose, human blood, human bone marrow, heart, kidney, lung, skeletal muscle, skin, and spleen, **Supplementary Figure S4**).

### Primate clocks

We developed two pan-primate clocks that are expected to apply to all primate species. Toward this end, we used all available primate tissues (**Supplementary Table S1**). The two primate clocks estimate chronological age and relative age, respectively. Relative age is defined as the ratio of chronological age to the respective maximum lifespan of the species (**Supplementary Table S3)** based on an updated version of the “anAge” data base [23].

To evaluate the primate clocks, we used a 10-fold cross validation scheme that was balanced with respect to species, i.e. each fold contained the same proportion of species. We found that a square root transformation of age, more precisely sqrt(Age+1), led to superior performance compared to the more widely used logarithmic transformation or identity transformation. The offset term of 1 year in the age transformation ensures that the clock is applicable to fetal samples as well (which are coded using negative age values). The primate clock for chronological age was highly accurate (R=0.99, **Figure 2A**) across all 37 primate species with an age correlation exceeding 0.90 in each species (**Figure 2B-F**). The lowest accuracy was observed for blood and skin samples from strepsirrhini species (R=0.90, **Figure 2I**), which probably reflects that the underlying data come from 26 strepsirrhini species that diverged many millions of years ago, e.g., *Varecia rubra* diverged from *Eulemur fulvus* about 24.8 million years ago [21]. The primate clock for age also performs well with humans (R=0.98, median error=2.11 years, **Figure 2E**) and compares favorably to the most accurate human clocks [24]. The primate clock for *relative* age is less accurate than that for chronological age (R=0.96 versus R=0.99, **Supplementary Figure S5**) but it may be attractive for specific applications that want to adjust for the substantial differences in maximum lifespan. Details on the CpGs underlying the clocks can be found in **Supplementary Table S4**.

**Figure 2:**
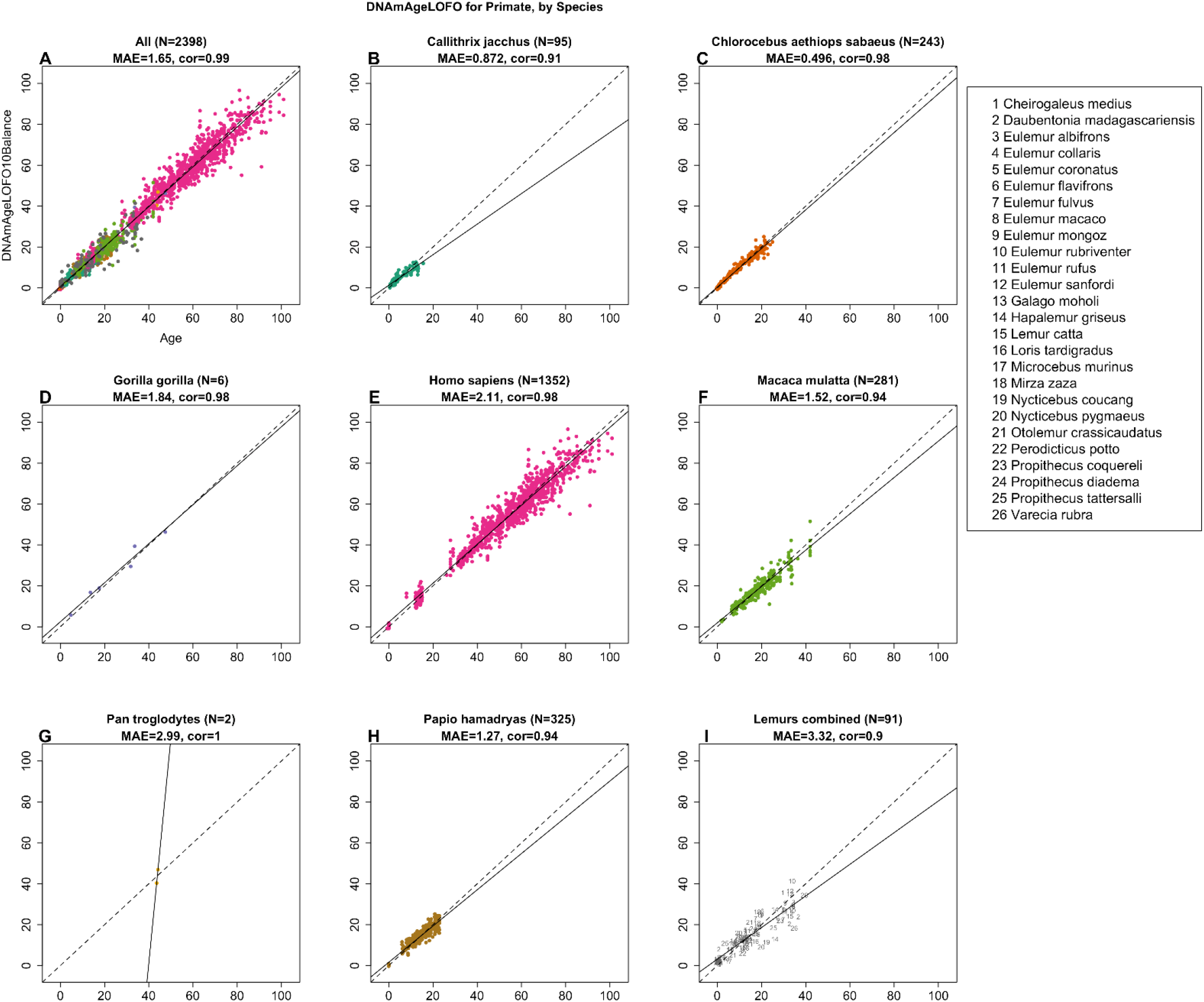
Cross-validation study of the primate clock for chronological age. A) Epigenetic clock based on tissues from all primate species (colored by species as indicated in the other panels). B-I) are excerpts from panel A but restricted to specific species mentioned in the title. I) Samples (dots) are blood and skin samples from 26 species of strepsirrhines as indicated in legend. Chronological age (years, x-axis) versus the ten-fold cross validation (balanced by species) of age (y-axis). Each panel reports the sample size, correlation coefficient, median absolute error (MAE). Dots are colored by species. The primate clock was developed by regressing sqrt(Age+1) on cytosines that map to baboons and humans. We also developed a primate clock for relative age, defined as the ratio of age by the respective maximum lifespan (Supplementary Figure S8). Details can be found in the Supplement.

### EWAS of age in baboons

To study tissue and species specific aging effects on cytosine levels, we carried out an Epigenome-Wide Association study (EWAS) of age in different baboon tissues (**Figure 3A**). The EWAS results for other primate species can be found in our companion papers [18, 19] (Methods). Overall, EWAS results in one baboon tissue correlate weakly with those in another tissue (**Supplementary Figure S6**) similar to what has been observed in humans [25]. However, the low conservation of aging effects across tissue types may reflect a limited sample size of non-cortex tissues (**Supplementary Table S2**).

**Figure 3.**
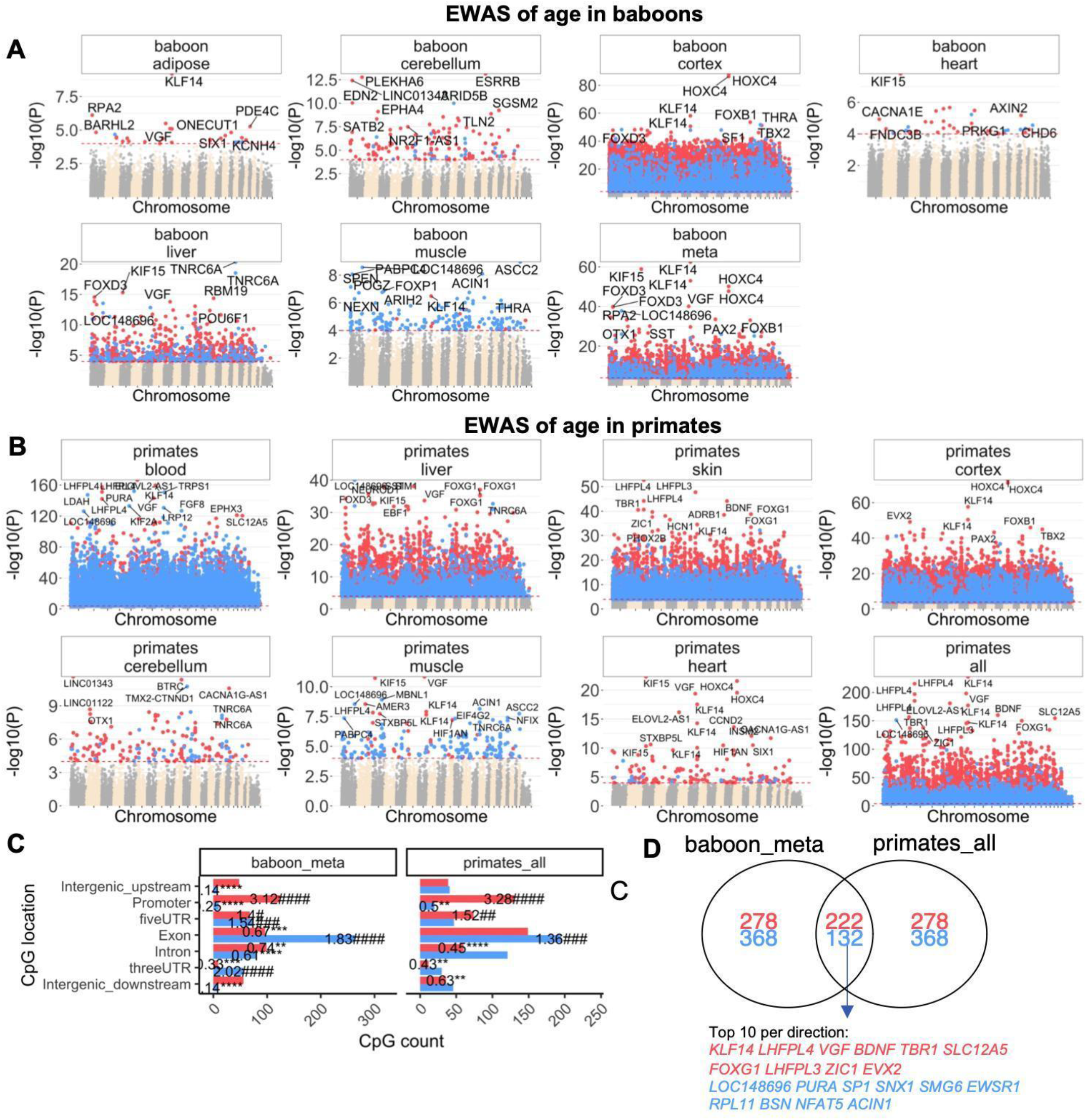
EWAS of age in baboons and primates. Meta-analysis of correlation tests in. A) Manhattan plots of the EWAS of chronological age in different baboon tissues (adipose, cerebellum, cerebral cortex, heart, liver, muscle of olive baboons). Note that the cerebral cortex leads to more significant p values (-log base 10 transformation of the p value, y-values) which reflects the broader age range of this tissue. B) Manhattan plots of the EWAS of chronological age primates. The p-values are based on Stouffer Z score meta-analysis of correlation with age in different tissues of primate species. “Primate-all” group is the Stouffer meta-analysis of age across all tissue types. The genome coordinates are estimated based on the alignment of Mammalian array probes to the Human Hg19 assembly. The direction of associations with p < 10^−4^ (red dotted line) is color-coded in red (age related increase in methylation) or blue (age related decrease). The top 15 CpGs were labeled by neighboring genes. C) Location of top CpGs (up to 500 per direction at p < 10^−4^ significance) in each tissue relative to the closest transcriptional start site. Significant odds ratios (count versus background) are reported for each bar. Fisher exact p values are encoded as follows *,# p<0.05, **,## p<0.01, ***,###p<0.001, ****,####p<0.0001. D) Venn diagram of the overlap of the meta-analysis of age in baboons and primates.

To capture the most-affected loci in all baboon tissues, a nominal p value < 10^−4^ was employed. Apart from p values, we also report correlation test Z statistics, which follow a standard normal distribution under the null hypothesis of zero correlation. Positive and negative values of the Z statistics correspond to age related gain/loss of methylation. The top age related CpGs in specific baboon tissues were located in the following genes: *KLF14* promoter (Z=6.2, adipose); *ESRRB* intron (Z=7.5, cerebellum); *HOXC4* promoter (Z=19.9, cerebral cortex); *KIF15* promoter (Z=5.6, heart); *TNRC6A* exon (Z=-9.4, liver); and *ASCC2* exon (Z=-6.1, muscle). To identify age-associated CpGs that are shared across the tissues, we carried out Stouffer’s meta-analysis across the six tissue types. Meta analysis across baboon tissues implicated *KLF14* (Z=16.7), *KIF15* (Z=16.3), and *HOXC4* (Z=15). In addition, two CpGs in the exon of *FOXD3* (Z=13.3) exhibit positive correlations with age (**Supplementary Table S5**). Not all of these CpGs correlate with age in strepsirrhines as shown in the following.

### EWAS of age in all primates

For seven different tissue types (Blood, Cerebellum, Cerebral Cortex, Heart, Liver, Skeletal Muscle, Skin), we carried out a meta analysis EWAS of age across primate species (**Figure 3B**). We observe tissue specific aging patterns (**Supplementary Figure S6**). For example, the top EWAS hits in primate blood (*LHFPL4, KLF14*) are not among the top hits in primate liver (**Figure 3B**). When focusing on blood samples, *ELOVL2-AS1* is particularly noteworthy since *ELOVL2* has been implicated in human epidemiological studies [4, 26]. Cytosine methylation levels in *ELOVL2* appear to regulate aging in the retina [27].

To identify CpGs that relate to age in most tissues and primates, we used a two-step meta analysis approach to combine the EWAS results of age across 29 strata of primate species and tissue type for which at least 10 samples were available. We find that CpGs near *KLF14* and *LHFPL4* gain methylation in most primate tissues (**Supplementary Table S6)**. A CpG near *ELOVL2-AS1* (cg16867657) correlates positively with age in several species (*Chlorocebus aethiops sabaeus, Homo sapiens, Macaca mulatta*, and *Papio hamadryas)* but not in *Callithrix jacchus* or strepsirrhines (**Supplementary Table S6**). A CpG (cg14361627) underlying *KLF14* is significantly correlated with age in all Haplorhine species but not in strepsirrhines (**Supplementary Table S6**).

Many CpGs correlate *positively* with age in all considered primate species (both haplorhines and strepsirrhines) including three CpGs near *LHFPL4* (cg12841266, cg24866418, cg11084334) and CpGs near *LHFPL3, BDNF, TBR1, SLC12A5, FOXG1* (**Supplementary Table S6**).

Conversely, we find CpGs that correlate *negatively* with age in all primate species including CpGs near *TRPS1 SNX1, SMG6, ARID5B, EWSR1* (**Supplementary Table S6**). However, we caution the reader that CpGs near these genes can have tissue specific aging patterns.

There is strong agreement between our EWAS findings in baboons and those across all primates. Out of the top 500 CpGs with a positive age correlation in baboon tissues, 222 overlap with the top 500 CpGs with a positive age correlation across all primate tissues (**Figure 3D**). A lower overlap (132 out of 500 CpGs) between baboons and primates can be observed for CpGs with negative age correlations. Overall, we find that CpGs with negative age correlations exhibit lower conservation across tissue type and primate species (**Figure 3D**).

Age-related CpGs map to both genic as well as intergenic regions, which are defined relative to transcriptional start and stop sites (**Figure 3C**). However, the relationship to age seems to depend on the relative gene region of the CpG. CpGs in promoters tend to gain methylation with age in primates, including baboon samples (odds ratio>3.3, p<10^−4^; **Figure 3C**). In contrast, the CpGs in exon regions tend to loose methylation with age (odds ratio>1.3, p<0.01; **Figure 3C**).

Enrichment studies show that CpGs with positive age correlations tend to be located i) in regions targeted by Polycomb Repressive Complex 2 (characterized by histone mark H3K27ME3) in embryonic stem cells, ii) near genes that play a role in development, and iii) in regions bound by transcription factors CHX10 and FOXO4 (**Figure 4**). CpGs with negative age correlations are located near genes that play a role in mRNA processing (**Figure 4, Supplementary Table S7**).

**Figure 4.**
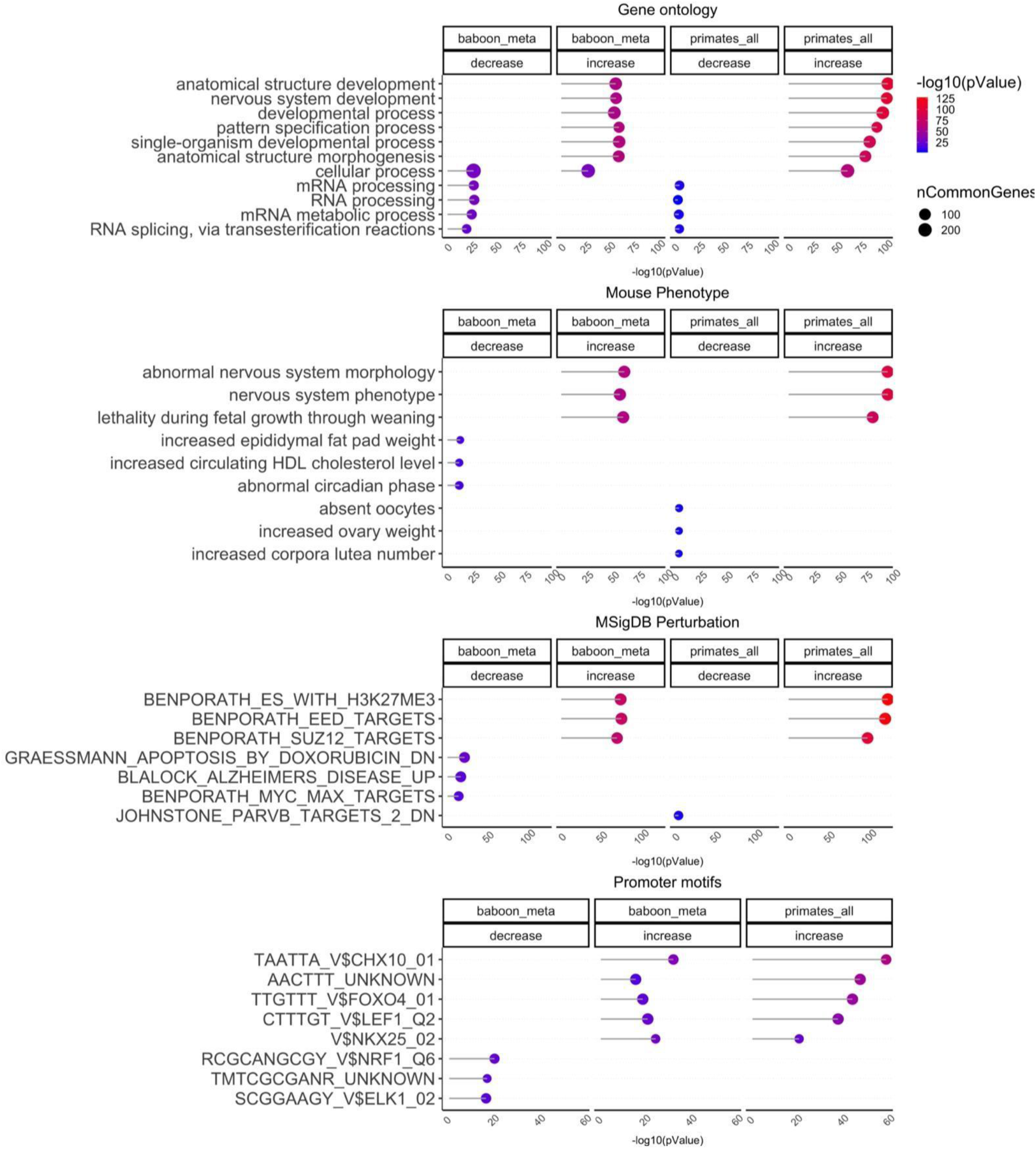
Gene set enrichment analysis of aging effects in primates. The gene level enrichment was done using GREAT analysis [52] relative to the array background and the human Hg19 background. The CpGs are annotated with adjacent genes within 50kb. We extracted up to top 500 CpGs based on p value of association per direction of change as input for the enrichment analysis. The p values are calculated by hypergeometric test of the EWAS results with the genes in each background dataset. Datasets: gene ontology, mouse phenotypes, promoter motifs, and MsigDB Perturbation, which includes the expression signatures of genetic perturbations curated in GSEA database. The results were filtered for significance at p < 10^−5^ and only the top terms for each EWAS are reported.

### Sex effects in primates

We aimed to find CpGs that relate to sex in many different tissues and primate species. However, we excluded the common marmoset from our analysis, because marmosets appear to have a unique characteristic when it comes to DNA methylation studies in mammals: it is not possible to predict sex on the basis of methylation data in common marmosets [19]. This curious observation probably reflects that the blood from marmosets is chimeric. Marmosets nearly always produce pregnancies consisting of at least dizygotic twins and during gestation fetuses exchange hematopoietic stem cells, which are not rejected. As a result, nearly all newborn, juvenile and adult marmosets carry a proportion of blood cells that do not reflect their own germline inheritance, but rather the germline of its dizygotic co-twin [28]. And because this process occurs with opposite sex dizygotic twins, it is common for female marmosets to carry blood cells derived from their brothers (with Y-chromosomes) and males to carry blood cells from their sisters.

We used the data from the other primates to identify sex-related CpGs in multiple tissues. For each species, we focused on the subset of CpGs that map to the respective genome (e.g. 37492 CpGs for humans, 34939 for vervet monkey, 34865 for baboons). To find sex related CpGs, we regressed cytosine methylation (dependent variable) on both age and sex. In total, the analysis involved 36 regression models in 13 tissues from human, baboon, vervet monkey, macaque, and strepshirrhine primates from prenatal time to late adulthood.

The median number of sex-associated methylation positions (SMP) in all these models was 1117 CpGs (at a 5% False Discovery Rate), most of which were located on the X chromosome. The largest number of SMPs (more than 7000 CpGs) could be found blood samples from macaque and vervets aged above 1 (**Figure 5A**). By contrast, we found only 1022 SMPs in human blood. In humans, the heart and muscle samples had a higher number of SMPs (2497 and 2226 CpGs, respectively). In specific tissues (heart and muscle in humans, blood in macaque and vervet monkey), we find that a surprisingly large number of SMPs are located on *autosomes* but in most tissues, the number of autosomal SMPs is very limited (fewer than 200 in most tissues). In total, 780 SMPs were significantly associated with sex in more than 19 model out of 36 regression models. 25 X chromosomal CpGs are related to sex in 32 models (**Supplementary Table S8**).

**Figure 5.**
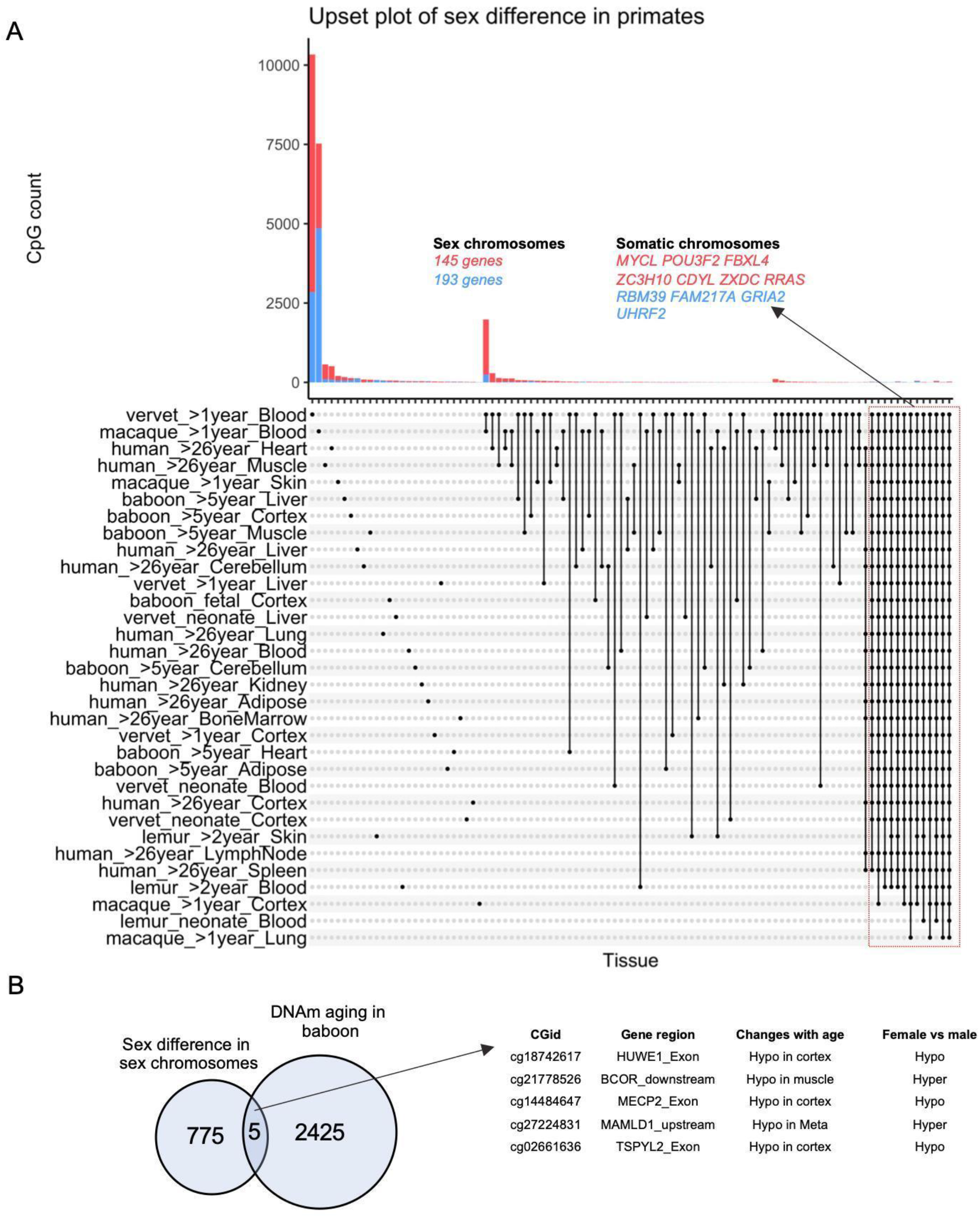
Sex effects on DNA methylation in primates. A) The upset plot of the sex-mediated differential methylated CpGs in 13 tissues from 30 primate species (human, baboon, vervet monkey, macaque, and 26 strepsirrhine primate species which where were analyzed together and denoted as lemur) at different age ranges. Red (blue) color codes genes/CpGs with significantly increased (decreased) methylation levels in females compared to males. The analysis included 30 multivariate regression models (stratified by species, tissues, and age categories), including chronological age as the covariate. The y-axis reports the number of CpGs that are significant at a 5% FDR rate. Neonates in vervet monkeys and strepsirrhine primates were aged <1 year. 780 CpGs adjacent to 248 genes related to sex in >19 of these models. B) The overlap of conserved sex-dependent CpGs and DNAm aging in the baboons. Manhattan plots of the sex differences in specific baboon tissues can be found in Supplementary Figure S9. The box plot of sex differences in these CpGs are reported in Supplementary Figure S10.

As expected, most of the sex-related CpGs, namely 769 CpGs, are located on sex chromosomes reflecting X chromosomal inactivation in females. The 11 autosomal sex-related CpGs (1.3%) are proximal to 11 genes (**Figure 5A, Supplementary Table S8**). Three of these genes (*FAM217A, CDYL, POU3F2*) are located in human chromosome 6 while the remaining autosomal sex-related CpGs are located near *MYCL, ZC3H10, ZXDC, RRAS, RBM39, GRIA2, UHRF2*, and *FBXL4* (**Figure 5A**).

Of the 780 SMPs, only 5 CpGs correlate significantly with age in baboons (CpGs near *HUWE1, BCOR, MECP2, MAMLD1, TSPYL2*, **Figure 5B**).

Detailed enrichment analysis of sex-differences in all primates can be found in **Supplementary Figure S7** and **Supplementary Table S9**. As expected, enrichment analysis of the sex-associated CpGs highlighted gonosomal and X-linked inheritance for CpGs located on both sex or autosomal chromosomes in all primates. The genes on sex chromosomes were also related to testicular development and several cognitive and neuronal pathways including intellectual disability, cognitive impairment, autism spectrum disorder, mental health, agitation, and thermal nociceptive threshold. Strikingly, the genes on the autosomal chromosome were also enriched in several neuronal pathways such as GABA receptor activation, intellectual disability, and synaptic function. Thus, our result suggests a possible difference in neuronal development, activity or function between sexes across these primate species that are associated with epigenetic changes. These differences are apparent from the fetal stage to very old age.

### Sex effects in baboons

We used the baboon samples to provide a detailed analysis of sex effects in different tissues and age groups. These results largely replicate the above mentioned findings for all primates. For example, the same 11 autosomal locations continue to be associated with sex (**Supplementary Figure S8, Supplementary Table S8**). The GREAT enrichment analysis in baboons leads to similar enrichments including X-linked inheritance and intellectual disability (**Supplementary Figure S9**).

### Sex predictors in all primates

Using elastic net regression, we developed a multivariate predictor of sex that applies to all primates except for marmosets (**Supplementary Table S10**). The predictor of female sex is based on 86 CpGs that are located on the X chromosome. A cross validation analysis suggests that the predictor is 99% accurate, e.g. it misclassified a single human sample (out of n=1352 samples). We also carried out an EWAS of sex that ignored species and tissue type. Across all primate samples from 37 primate species, 294 X-chromosomal CpGs markers were significantly hypomethylated and 165 CpGs were hypermethylated in females at a significance threshold of p=10^−300^ (**Supplementary Table S11**). Boxplots for six representative CpGs are presented in **Supplementary Figure S10**.

### Sex effects in large human cohorts

To test whether the sex-specific results in primates could be corroborated in human methylation data generated on another genomic platform, we analyzed human Illumina EPIC array data from three large studies: blood samples from individuals of European ancestry (Framingham Heart study), blood samples from individuals of African ancestry (Jackson Heart study), and postmortem prefrontal cortex samples from individuals from the Religious Order study (ROSMAP). CpGs can fall in several regions relative to the transcriptional start site of a gene e.g. the gene promoter, gene exon, 5’ untranslated regions, 3’ UTR, intergenic regions.

Since the mammalian array and the EPIC array share only 7111 CpGs [22], we compared the results of the two platforms at the level of gene region. Almost all human gene-regions that contain at least one CpG from the mammalian array are also covered by the human EPIC array (**Supplementary Figure S11**). Several gene-regions proximal to these SMPs in primates were also differentially methylated between human males and females in both blood and brain samples (**Figure 6**). Interestingly, this analysis also highlighted autosomal gene *POU3F2* on chromosome 6, which was one of the 11 autosomal sex-related genes across primates. Significant sex-related CpGs in human brain samples are reported in **Supplementary Figure S12**.

**Figure 6.**
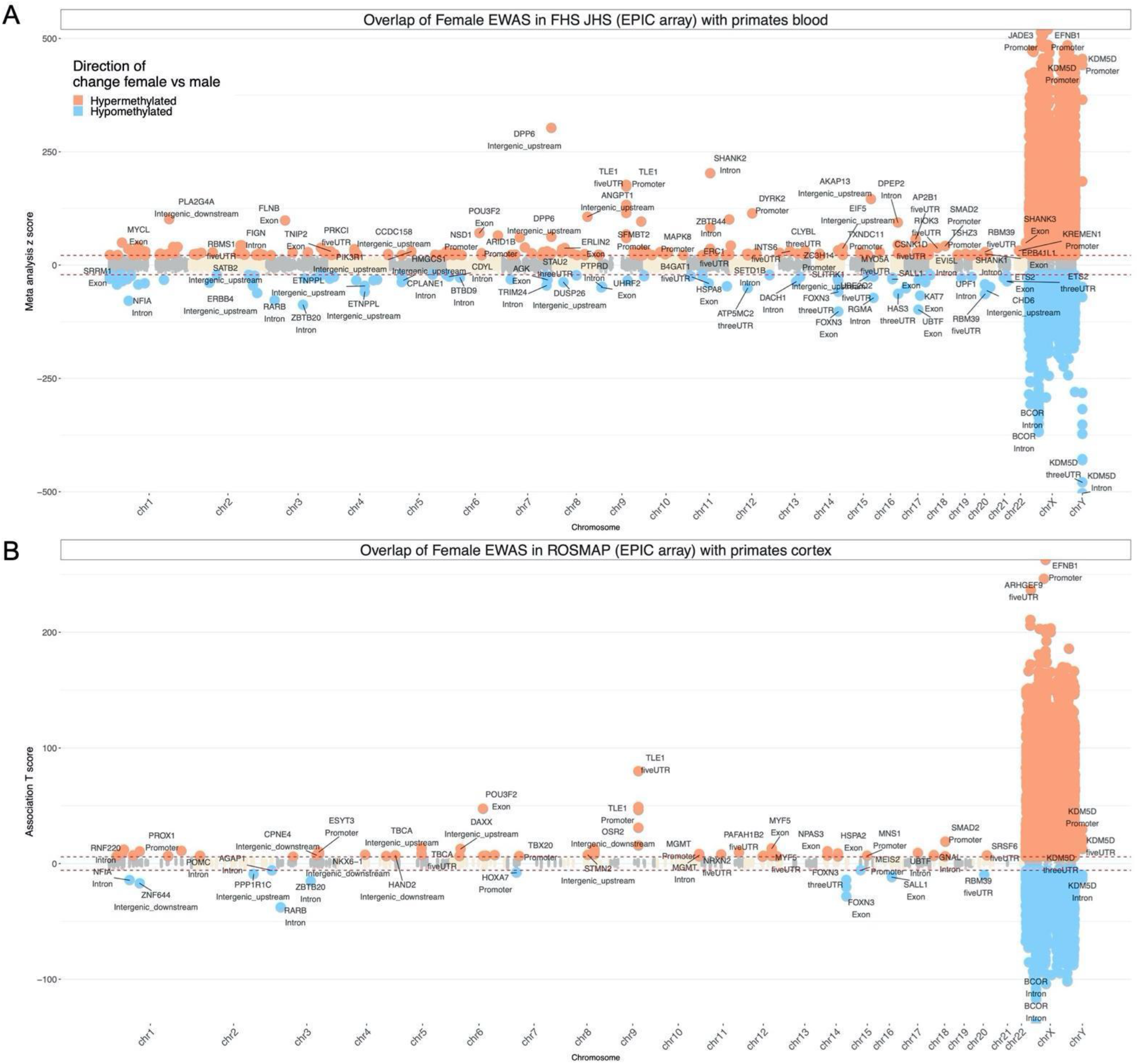
Sex related methylation changes in primates implicated by two array platforms. The Manhattan plots only involves EPIC array probes that map to the same human gene regions as the corresponding probe on the mammalian array. The y-axis reports sex effects (Z statistics) based on the human EPIC array. The x-axis reports genomic locations (gene regions) that were significant (5% FDR) in at least one of the primates blood samples (human, macaque, vervet monkey). A) The overlap of primates EWAS of sex with the meta-analysis of blood samples from the Jackson Heart (JHS) and Framingham Heart (FHS) Studies. Red (blue) color codes genes/CpGs with significantly increased (decreased) methylation levels in females compared to males. The meta-analysis z score is calculated by Stouffer’s method from the EWAS of sex (adjusted for age) in JHS and FHS. The red lines are z scores that corresponds to p<10^−100^ in EPIC array meta-analysis. The top two gene regions from each chromosome were labeled. B) The overlap of primates EWAS of sex (adjusted for age) in prefrontal cortex samples from ROSMAP. The probes were also further filtered for the gene regions that were significant (5% FDR) in at least one of the primates’ cerebral cortex samples (baboon, vervet monkey). The red lines are Student T test statistics that corresponds to a significance threshold of p<10^−8^ in the EWAS analysis. The human EPIC array covers most of the gene regions that are covered by the mammalian methylation array (**Supplementary Figure S11**).

## Discussion

Apart from two primate clocks that apply to all primate species, we present five epigenetic clocks that are tailor-made for baboons and are applicable to their entire life course (from birth to old age). Our genomic platform (mammalian array) focused on highly conserved CpGs which allowed us to develop epigenetic clocks that are compatible with different primate species even though humans separated from the old-world monkey lineage approximately 23-25 million years ago, the baboon and rhesus macaque lines separated approximately 9 million years ago, and the baboon and vervet monkeys separated 10 million years ago [21]. Lemurs are an evolutionary puzzle since their island home (Madagascar) separated from Africa about 145 million years ago, before the origin of placental mammals. Fossil evidence however, suggests that lemurs evolved closer to 20 million years ago.

Despite the substantial evolutionary distance between the 37 primate species, we managed to construct two highly accurate primate clocks based on a single multivariate model. This effectively consolidates the notion that certain epigenetic aspects of aging are highly conserved between primate species. Interestingly, our primate clock for chronological age is more accurate than the clock for relative age. However, it was important to apply a square root transformation to age to achieve this high accuracy.

The ability to generate primate clocks should not detract from the fact that there are profound differences in age-related methylation changes across species and even between tissues from the same species. Indeed, when age-associated CpGs were analyzed, their tissue specificity was readily apparent (**Supplementary Figure S6**). This can appear at first sight to be inconsistent with the successful development of a pan-tissue clock. It is however important to bear in mind that age-associated CpGs that constitute the clock are not necessarily the highest scoring ones for individual tissues. As such, it appears that as tissues age, two sets of CpGs undergo methylation change; the first is tissue-specific and the second is general across all tissues, which is consistent with what has been observed in humans [3, 29, 30].

A CpG in the promoter of *KLF14* was the top age-related CpG across all baboon tissues and also the top hit in our meta-analysis of primate tissues (**Figure 3**) but this CpG does not correlate with age in the strepsirrhines (**Supplementary Table S6**). *KLF14*, is a transcription regulator that can activate and also repress the expression of genes. Although its direct targets remain to be fully characterized, its pathologic effect is known to include prevention of cardiovascular disease [31]. Methylation of *KLF14* locus in blood of mice is predictive of chronic inflammation in adipose tissue [32], suggesting that this cross-tissue age-associated locus might extend to other tissues and may bear even greater predictive scope. Indeed, *KLF14* is also one of the top EWAS findings in primate blood. Human epidemiological studies have revealed that methylation of this locus in blood, saliva and buccal swabs is predictive of age in humans [33], and along with *ELOVL2, KLF14* is one of the most consistently identified loci with age-associated methylation change [34].

We identified many CpGs that showed a consistent aging pattern across all primate species (haplorhines and strepsirrhines) e.g. the positively age-related CpGs near *LHFPL4, LHFPL3, BDNF, TBR1, SLC12A5, FOXG1* and negatively age-related CpGs near *TRPS1, SNX1, SMG6, ARID5B, EWSR1* (**Supplementary Table S6**). Our meta-analysis of age across baboons and primate tissues implicated CpGs near several members of Hox family genes (e.g. *HOXC4, HOXA11, HOXD8*, and *HOXC10)* which gain methylation with age. The Hox family is one of the oldest gene sets that are conserved across not only mammals but arthropods as well. Hox proteins specify development of body segments of organisms and are pivotal for normal healthy development. Their expression is regulated by polycomb repressive complex, whose DNA targets are consistently identified, as well in this study, as being increasingly methylated with age [3, 7]. Hence the involvement of HOX loci further implicates the importance of the developmental process in aging. This is further validated through the enrichment analyses of age-associated CpGs that were identified with developmental processes across the different tissues.

The specific baboons used to develop the reported clocks herein are hybrid animals resulting from the mating between olive and yellow baboons. While, we could not evaluate whether methylation rates might be different in hybrids as compared to pure-bred members of one or the other type. However, our experience with other species leads us to predict that any differences would be minor. In the worst case, the baboon clocks would be offset by a constant value between DNAmAge and actual age. By definition, our primate clock applies to all primate species and would certainly apply to different species of baboons.

Sex affects the blood methylome in a species and life stage dependent manner. In neonatal blood from vervets, we observed few autosomal CpGs that relate to sex but this number increases dramatically in older animals. While blood samples from older macaques and vervets also reveal a large number of autosomal CpGs that relate to sex, the same could not be observed in human blood samples. We present CpGs that relate to sex in all primate species. A subset of these CpGs allowed us to develop a multivariate estimator of sex that applies to all primate species and all tissue types. Sex predictors can be used to find plate map errors or other data handling errors. Most of the sex-related CpGs were located on the X chromosome. However, our EWAS of sex also identified 11 autosomal locations. Methylation array data generated on two different platforms (mammalian array and human EPIC array) highlight the autosomal sex-related gene *POU3F2* on human chromosome 6. Another autosomal sex-related CpG was located near *CDYL*, which encodes a positive regulator of polycomb repressive complex 2 (PRC2) [35, 36]. This is particularly relevant as PRC2 target sites become increasingly methylated with age across different mammalian species [3, 37]. The collective evidence from many different species analyzed thus far repeatedly points to PRC2 as a pivotal factor underlying epigenetic aging. Therefore, the difference in methylation rate of the *CDYL* locus could potentially result in differences in PRC2 activity.

Another group has previously presented a blood clock for baboons based on DNA methylation profiles that were generated using a different methylation profiling platform (Reduced Representation Bisulfite Sequencing) [38]. We could not evaluate this blood clock in our data because the two genomic platforms profiled different sets of CpGs and furthermore, we did not profile blood samples in baboons.

Beyond their utility, the primate and baboon epigenetic clocks reveal several salient features with regard to the biology of aging. First, the fact that these are pan-tissue clocks re-affirm the notion that aging is a coordinated biological process that is harmonized throughout the body. Second, the developmental process is further implicated as being an important component of the process of epigenetic aging. Third, the ability to develop human-baboon clocks and even primate clocks attests to the high conservation of the aging process across evolutionary distinct primate species. This implies that treatments that alter the epigenetic age of baboons, as measured using the human-baboon clock are very likely to exert similar effects in humans.

We expect that the availability of the baboon and primate clocks will provide a significant boost to the potential use of baboons and other primates as models for human aging.

## Materials and Methods

### Baboon care and maintenance

All animals were given a full veterinary examination prior to recruitment to the study and no obvious cause of ill health or pathology was observed. The animals were housed in group cages at the Southwest National Primate Research Center, at Texas Biomedical Research Institute (TBRI), in San Antonio, Texas in mixed sex groups of up to 16. The remaining 4 females were housed in individual cages at the UT Health Sciences Center San Antonio (UTHSCSA).

All procedures were approved by the TBRI or UTHSCSA Animal Care and Use Committee and conducted in facilities approved by the Association for Assessment and Accreditation of Laboratory Animal Care.

Twenty-eight females and the ten males were fed ad libitum Purina Monkey Diet 5038 (12% energy from fat, 0.29% from glucose and 0.32% from fructose and metabolizable energy content of 3.07 kcal/g protein; Purina LabDiets, St Louis, MO, USA) (CTR). Water was continuously available to all animals. Animal health status was recorded daily.

#### Necropsy

None of the animals were euthanized for this project. Rather, we used left-over frozen tissue samples that had previously been collected as part of other projects. Necropsies were performed by either a qualified, experienced veterinarian or M.D investigator. At TRBI, baboons were pre-medicated with ketamine hydrochloride (10 mg/kg IM) and anesthetized using isoflurane (2%) resulting in general anesthesia as previously described [39]. Baboons were exsanguinated while under general anesthesia as approved by the American Veterinary Medical Association. At UTHSCSA four animals were euthanized using Pentobarbital at 390 mg/ml (Fatal-Plus Solution, Vortech, Dearborn, MI, USA). Following cardiac asystole, respiratory failure and a lack of reflexive response to both skin pinch and eye touch stimulation, tissues (adipose, cerebellum, cerebral cortex, muscle, heart, liver) were rapidly dissected and immediately frozen in liquid nitrogen.

For the studies in which fetal tissue was obtained, all animals were housed in 20 foot × 20 foot × 15 foot metal and concrete group cages at the Texas Biomedical Research Institute. Experimental animals were obtained from appropriate groups of 16 healthy female baboons of similar pre-study body weights (10–15 kg) and morphometric features (13). The potential day of conception was determined based on the day of ovulation and changes in sex skin color and pregnancy was confirmed at 30 days post ovulation by using ultrasonography. Details of housing, feeding, and environmental enrichment have been published elsewhere (13). All procedures were approved by the University of Texas Health Science Center and Texas Biomedical Research Institute internal animal care and use committees and performed in the Association for Assessment and Accreditation of Laboratory Animal Care–approved facilities.

Prior to Cesarean section, baboons were premedicated with ketamine hydrochloride (10 mg/kg, IM). Following tracheal intubation, isoflurane (2%, 2L/min, by inhalation) was used to maintain an appropriate plane of anesthesia throughout the surgery. A cesarean section was performed at gestational day 165 (0.9 of gestation) using standard sterile techniques as previously described [40]. Following hysterotomy, the umbilical cord was identified and used for fetal exsanguination with both maternal and fetal baboon under general anesthesia as approved by the American Veterinary Medical Association Panel on Euthanasia. Postoperatively, mothers were placed in individual cages and watched until they were upright under their own power. Maternal analgesia was administered for 3 days (buprenorphine hydrochloride injection; Hospira, Inc., Lake Forest, IL, USA; 0.015 mg/kg/day) post-operatively or longer if indicated. They were returned to their group cage two weeks postoperatively.

Animals were individually fed to enable precise regulation of intake either between 7:00 am and 9:00 am or 11:00 am and 1:00 pm as described in detail elsewhere [41]. Water was continuously available in each feeding cage (Lixit, Napa, California), and the animals were fed Purina Monkey Diet 5038 (Purina, St Louis, Missouri). For this study, we selected samples representing the entire primate lifespan, from neonate to old age (**Supplementary Table S2**).

#### Strepsirrhine primates

Strepsirrhini is a suborder of primates that includes the lemuriform primates, which consist of the lemurs of Madagascar, pottos and galagos from Africa, and the lorises from Southeast Asia. Lemuroids and lorisoids together form the more ancestral sister clade to all other living primates. As such, they lend unparalleled power to any comparative study within the primate clade, or to studies comparing primate features to those of other mammals.

For this study, we selected a total of 91 samples from individuals representing 26 strepsirrhine species, in most cases, the entire lifespan, from immature (infant or juvenile) to senile stages: 68 samples from peripheral blood, 23 samples from skin (**Supplementary Table S1**). The strepsirrhine primates (suborders Lemuriformes and Lorisiformes) used in this study were from the Duke Lemur Center (DLC) in Durham, NC (USA). The Duke Lemur Center is certified by both the Association for Assessment and Accreditation of Laboratory Animal Care and the American Zoological Association. The animal handling and sample collection procedures in this study were performed by a veterinarian after review and approval by the Duke Institutional Animal Care and Use Committee and the DLC Research Committee. Both housing and sample collection met or exceeded all standards of the Public Health Service’s “Policy on the Humane Care and Use of Laboratory Animals”.

The lemurs are housed in comparable social and housing conditions, habituated to human presence, and individually identifiable. The DLC also maintains a large collection of banked tissues, deriving from routine veterinary procedures and necropsies, amassed over the Center’s 55-year history. These samples represent a broad diversity of taxa (>30), individuals (>4,500), and specimen type (e.g., blood, organ tissues, cadavers). Detailed records of life and medical history, reproduction, and social-group membership are digitally maintained [42].

The DLC makes available its collection of live animals, cadavers, tissues, and microCT-imaged skeletons for research and teaching. The resulting projects span a broad array of biological disciplines including, but not limited to, behavioral ecology, biomechanics, cognition, anatomy, life-history strategy, metagenomics, molecular evolution, phylogenomics, population genetics, metabolomics, sensory communication, and speciation. The DLC maintains a policy of “do no harm” research that is compatible with the critically endangered status of lemurs and other strepsirrhines. While we are dedicated to pushing the frontiers of minimally-invasive research, augmented by emerging technologies such as genomics and proteomics, we are committed to maintaining and sustaining a healthy and reproductively active colony. Moreover, research policies ensure that the data collected are unbiased by stress or compromised animal health.

Peripheral blood was collected through venipuncture with standard procedures, either during a routine veterinary procedure or at time of necropsy. Skin tissues were collected during necropsies. Whole blood was preserved in either EDTA or Lithium Heparin, and stored at −80oC. Skin tissues were either frozen directly at −80oC or were first flash frozen and then stored at −80oC.

We profiled the following species: Cheirogaleus medius (Fat-tailed dwarf lemur), Daubentonia madagascariensis (Aye-aye), Eulemur albifrons (White-headed lemur), Eulemur collaris (Collared brown lemur), Eulemur coronatus (Crowned lemur), Eulemur flavifrons (Blue-eyed black lemur), Eulemur fulvus (Brown lemur), Eulemur macaco (Black lemur), Eulemur mongoz (Mongoose lemur), Eulemur rubriventer (Red-bellied lemur), Eulemur rufus (Red-fronted lemur), Eulemur sanfordi (Sanford’s brown lemur), Galago moholi (South African galago), Hapalemur griseus (Bamboo lemur), Lemur catta (Ring-tailed lemur), Loris tardigradus (Slender loris), Microcebus murinus (Gray mouse lemur), Mirza zaza (Northern giant mouse lemur), Nycticebus coucang (Slow loris), Otolemur crassicaudatus (Greater galago), Perodicticus potto (Potto), Propithecus diadema (Diademed sifaka), Propithecus tattersalli (Golden-crowned sifaka), Varecia rubra (Red ruffed lemur).

#### Primates from Busch Gardens

The blood samples from chimpanzees (*Pan troglodytes*, n=2), gorillas (*Gorilla*, n=3)

Orangutan (n=1, *Pongo pygmaeus*), red ruffed lemur (n=1, *Varecia variegata*), and White-fronted marmoset (n=1, *Callithrix geoffroyi*) were opportunistically collected and banked during routine health exams from these zoo-based animals located at Busch Gardens Tampa (Tampa, Florida).

#### Existing data from primates

The data from other monkeys are described in the companion papers for rhesus macaque (skin, blood, adipose, cerebral cortex, liver, lung, muscle [18]), vervet monkey (whole blood, prefrontal cortex and liver [17]), and common marmosets (n=95 blood samples [19]).

#### Human tissue samples

We analyzed previously generated methylation data from n=1352 human tissue samples (adipose, blood, bone marrow, dermis, epidermis, heart, keratinocytes, fibroblasts, kidney, liver, lung, lymph node, muscle, pituitary, skin, spleen) from individuals whose ages ranged from 0 to 93 years. The tissue samples came from four sources: tissue and organ samples from the National NeuroAIDS Tissue Consortium [43], Blood samples from the Cape Town Adolescent Antiretroviral Cohort study [44] and the PEG study [45], skin and other primary cells provided by Ken Raj [46]. Ethics approval (IRB#18-000315).

#### DNA extraction

DNA was extracted on an automated nucleic acid extraction platform Anaprep (Biochain) using a magnetic bead based extraction method and Tissue DNA Extraction Kit (AnaPrep).

#### DNA methylation data

All methylation data were generated using a custom mammalian methylation array (HorvathMammalMethylChip40) based on 37492 CpG sites [22]. While all of these CpGs can be mapped to the human genome, not all of them can be mapped to other primates. We only used mappable CpGs in species specific analyses. For example, 34,865 CpGs from the mammalian array could be aligned to specific loci that are proximal to 5781 genes in the Olive baboon (Papio anubis.Panu_3.0.100) genome. The genome coordinates of each CpG probe on the mammalian array is provided on the Github page of the Mammalian Methylation Consortium https://github.com/shorvath/MammalianMethylationConsortium/

The manifest file of the mammalian array can also be found on Gene Expression Omnibus (GPL28271). The SeSaMe normalization method was used to define beta values for each probe [47].

#### Penalized Regression models

Details on the clocks (CpGs, genome coordinates) and R software code are provided in the Supplement. Penalized regression models were created with glmnet [48]. We investigated models produced by both “elastic net” regression (alpha=0.5). The optimal penalty parameters in all cases were determined automatically by using a 10-fold internal cross-validation (cv.glmnet) on the training set. By definition, the alpha value for the elastic net regression was set to 0.5 (midpoint between Ridge and Lasso type regression) and was not optimized for model performance.

We performed a cross-validation scheme for arriving at unbiased (or at least less biased) estimates of the accuracy of the different DNAm based age estimators. One type consisted of leaving out a single sample (LOOCV) from the regression, predicting an age for that sample, and iterating over all samples. A critical step is the transformation of chronological age (the dependent variable). While no transformation was used for the pan-tissue clock for baboons, we used a log linear transformation for the dual species clock of absolute age. For the primate clock, we used the following transformation: sqrt(Age+1), i.e. after adding an offset of 1 year, we formed the square root transformation. The square root transformation outperformed the log transformation and the identity transformation with respect to cross validation based estimates of accuracy.

We defined relative age as ratio: Relative age=Age/maxLifespan where the maximum lifespan for the different species was chosen from the anAge data base [23], e.g. the human maximum lifespan was determined to be 122.5. For the sake of reproducibility, we report the maximum lifespan and average age at sexual maturity in **Supplementary Table S2**.

#### Epigenome wide association studies of age in baboons

EWAS was performed in each tissue separately using the R function “standardScreeningNumericTrait” from the “WGCNA” R package. Next the results were combined across tissues using Stouffer’s meta-analysis method.

#### EWAS of age across all primate species

Our meta analysis for EWAS of age in primate species combined correlation test statistics calculated in 29 different species-tissue strata with a minimal sample size of 10 (N≥10). In the first stage, we combined the EWAS results across tissues within the same species to form species specific meta-EWAS results. In the second stage, we combined species EWAS results to form a final meta-EWAS of age. All the meta analyses in both stages were performed by the unweighted Stouffer’s method.

### Human studies on the EPIC array

#### Framingham Heart Study

We used blood methylation data from 2,356 individuals composed of 888 pedigrees from the Framingham Heart cohort [49], a large-scale longitudinal study started in 1948, initially investigating risk factors for cardiovascular disease (CVD). The FHS cohort contains blood DNA methylation profiling at exam 8. A human epigenetic clock analysis of these data is presented in many articles including [12, 50].

#### Jackson Heart Study (JHS, N=1747)

The JHS is a large, population-based observational study evaluating the etiology of cardiovascular, renal, and respiratory diseases among African Americans residing in the three counties (Hinds, Madison, and Rankin) that make up the Jackson, Mississippi metropolitan area. The age at enrollment for the unrelated cohort was 35-84 years; the family cohort included related individuals >21 years old. JHS ancillary study ASN0104, available with both phenotype and DNA methylation array data.

#### ROSMAP

We analyzed previously generated DNA methylation data from Caucasian subjects from the Religious Order Study (ROS) and the Rush Memory and Aging Project (MAP) [51] Both are longitudinal community based cohort studies of aging and dementia. The majority of participants in both studies are 75– 80 years old at baseline with no known dementia. All participants agree to organ donation at death. Participants sign and informed consent, repository consent, and Anatomical Gift Act.

## Acknowledgements

This work was mainly supported by the Paul G. Allen Frontiers Group (SH), 1U01AG060908-01 (SH), and Open Philanthropy (SH). We would like to acknowledge support through a 1U19AG057758-01A1 Womb to Tomb: Developmental programming and aging interactions in primates to PWN. This investigation used resources that were supported by the Southwest National Primate Research Center grant P51 OD011133 from the Office of Research Infrastructure Programs, National Institutes of Health.

We would like to thank the veterinary and technical staff at the Primate Center for their consistent help and advice on animal management. We would specifically like to express our gratitude to Ms. Susan Jenkins for maintaining the databases.

The rhesus macaque data were funded in part by the Intramural Research Program, National Institute on Aging, NIH. Blood samples from several primates were graciously provided by Busch Gardens Tampa.

The Jackson Heart Study (JHS) is supported by contracts HHSN268201300046C, HHSN268201300047C, HHSN268201300048C, HHSN268201300049C, HHSN268201300050C from the National Heart, Lung, and Blood Institute and the National Institute on Minority Health and Health Disparities.

The Framingham Heart Study is funded by National Institutes of Health contract N01-HC-25195 and HHSN268201500001I. The laboratory work for this investigation was funded by the Division of Intramural Research, National Heart, Lung, and Blood Institute, National Institutes of Health. The analytical component of this project was funded by the Division of Intramural Research, National Heart, Lung, and Blood Institute, and the Center for Information Technology, National Institutes of Health, Bethesda, MD. JMM and KLL were supported by R01AG029451.

The Religious Order study and Rush Memory and Aging Project were funded by P30AG10161, R01AG17917, RF1AG15819, R01AG34374, R01AG36042, U01AG46152 (Bennett).

This is Duke Lemur Center publication #1497. The vervet animal and tissue resources used here were supported by the following grants from the US National Institutes of Health: P40RR019963/ OD010965 (to M. J. Jorgensen); R01RR016300/OD010980 (to Nelson B.Freimer); R37MH060233 (to D. Geschwind). Marmoset resources were supported by funding from the National Institutes of Health through grants R01 AG050797, P30 AG013319, and P30 AG044271.

The funding bodies played no role in the design, the collection, analysis, or interpretation of the data.

## References

[1] H. Cedar and Y. Bergman, “Programming of DNA methylation patterns,” Annu Rev Biochem, vol. 81, pp. 97–117, 2012.

[2] V. K. Rakyan, T. A. Down, S. Maslau, T. Andrew, T. P. Yang, H. Beyan, et al., “Human aging-associated DNA hypermethylation occurs preferentially at bivalent chromatin domains,” Genome Res, vol. 20, pp. 434–9, Apr 2010.

[3] A. E. Teschendorff, U. Menon, A. Gentry-Maharaj, S. J. Ramus, D. J. Weisenberger, H. Shen, et al., “Age-dependent DNA methylation of genes that are suppressed in stem cells is a hallmark of cancer,” Genome Res, vol. 20, pp. 440–6, Apr 2010.

[4] P. Garagnani, M. G. Bacalini, C. Pirazzini, D. Gori, C. Giuliani, D. Mari, et al., “Methylation of ELOVL2 gene as a new epigenetic marker of age,” Aging Cell, vol. 11, pp. 1132–1134, 2012.

[5] I. Beerman, C. Bock, B. S. Garrison, Z. D. Smith, H. Gu, A. Meissner, et al., “Proliferation-dependent alterations of the DNA methylation landscape underlie hematopoietic stem cell aging,” Cell Stem Cell, vol. 12, pp. 413–25, Apr 4 2013.

[6] J. Przybilla, T. Rohlf, M. Loeffler, and J. Galle, “Understanding epigenetic changes in aging stem cells--a computational model approach,” Aging Cell, vol. 13, pp. 320–8, Apr 2014.

[7] M. Jung and G. P. Pfeifer, “Aging and DNA methylation,” BMC Biology, vol. 13, pp. 1–8, 2015.

[8] S. Horvath and K. Raj, “DNA methylation-based biomarkers and the epigenetic clock theory of ageing,” Nat Rev Genet, Apr 11 2018.

[9] C. G. Bell, R. Lowe, P. D. Adams, A. A. Baccarelli, S. Beck, J. T. Bell, et al., “DNA methylation aging clocks: challenges and recommendations,” Genome Biology, vol. 20, p. 249, 2019/11/25 2019.

[10] S. Horvath, “DNA methylation age of human tissues and cell types,” Genome Biol, vol. 14, p. R115, 2013.

[11] R. Marioni, S. Shah, A. McRae, B. Chen, E. Colicino, S. Harris, et al., “DNA methylation age of blood predicts all-cause mortality in later life,” Genome Biol., vol. 16, p. 25, 2015.

[12] B. H. Chen, R. E. Marioni, E. Colicino, M. J. Peters, C. K. Ward-Caviness, P. C. Tsai, et al., “DNA methylation-based measures of biological age: meta-analysis predicting time to death,” Aging (Albany NY), vol. 8, pp. 1844–1865, Sep 28 2016.

[13] M. E. Levine, A. T. Lu, A. Quach, B. H. Chen, T. L. Assimes, S. Bandinelli, et al., “An epigenetic biomarker of aging for lifespan and healthspan,” Aging (Albany NY), Apr 18 2018.

[14] A. T. Lu, A. Quach, J. G. Wilson, A. P. Reiner, A. Aviv, K. Raj, et al., “DNA methylation GrimAge strongly predicts lifespan and healthspan,” Aging (Albany NY), vol. 11, pp. 303–327, Jan 21 2019.

[15] L. A. Cox, A. G. Comuzzie, L. M. Havill, G. M. Karere, K. D. Spradling, M. C. Mahaney, et al., “Baboons as a model to study genetics and epigenetics of human disease,” ILAR journal, vol. 54, pp. 106–121, 2013 2013.

[16] A. M. Bronikowski, S. C. Alberts, J. Altmann, C. Packer, K. D. Carey, and M. Tatar, “The aging baboon: Comparative demography in a non-human primate,” Proceedings of the National Academy of Sciences, vol. 99, p. 9591, 2002.

[17] A. J. Jasinska, A. Haghani, J. A. Zoller, C. Z. Li, A. Arneson, J. Ernst, et al., “Epigenetic clock and methylation studies in vervet monkeys,” GeroScience, 2021/09/30 2021.

[18] S. Horvath, J. A. Zoller, A. Haghani, A. J. Jasinska, K. Raj, C. E. Breeze, et al., “Epigenetic clock and methylation studies in the rhesus macaque,” GeroScience, 2021/09/06 2021.

[19] S. Horvath, J. A. Zoller, A. Haghani, A. T. Lu, K. Raj, A. J. Jasinska, et al., “DNA methylation age analysis of rapamycin in common marmosets,” GeroScience, 2021/09/05 2021.

[20] L. Pozzi, J. A. Hodgson, A. S. Burrell, K. N. Sterner, R. L. Raaum, and T. R. Disotell, “Primate phylogenetic relationships and divergence dates inferred from complete mitochondrial genomes,” Molecular phylogenetics and evolution, vol. 75, pp. 165–183, 2014.

[21] S. Kumar, G. Stecher, M. Suleski, and S. B. Hedges, “TimeTree: A Resource for Timelines, Timetrees, and Divergence Times,” Molecular Biology and Evolution, vol. 34, pp. 1812–1819, 2017.

[22] A. Arneson, A. Haghani, M. J. Thompson, M. Pellegrini, S. B. Kwon, H. Vu, et al., “A mammalian methylation array for profiling methylation levels at conserved sequences,” bioRxiv, p. 2021.01.07.425637, 2021.

[23] J. P. de Magalhaes, J. Costa, and G. M. Church, “An analysis of the relationship between metabolism, developmental schedules, and longevity using phylogenetic independent contrasts,” J Gerontol A Biol Sci Med Sci, vol. 62, pp. 149–60, Feb 2007.

[24] S. Horvath, J. Oshima, G. Martin, K. Raj, and S. Matsuyama, “Epigenetic age estimator for skin and blood applied to Hutchinson Gilford Progeria,” 2018.

[25] T. Zhu, S. C. Zheng, D. S. Paul, S. Horvath, and A. E. Teschendorff, “Cell and tissue type independent age-associated DNA methylation changes are not rare but common,” Aging (Albany NY), vol. 10, p. 3541, 2018.

[26] M. G. Bacalini, J. Deelen, C. Pirazzini, M. De Cecco, C. Giuliani, C. Lanzarini, et al., “Systemic Age-Associated DNA Hypermethylation of ELOVL2 Gene: In Vivo and In Vitro Evidences of a Cell Replication Process,” The Journals of Gerontology: Series A, vol. 72, pp. 1015–1023, 2017.

[27] D. Chen, D. L. Chao, L. Rocha, M. Kolar, V. A. Nguyen Huu, M. Krawczyk, et al., “The lipid elongation enzyme ELOVL2 is a molecular regulator of aging in the retina,” Aging Cell, vol. 19, p. e13100, Feb 2020.

[28] C. Sweeney, J. Ward, and E. Vallender, “Naturally occurring, physiologically normal, primate chimeras,” Chimerism, vol. 3, pp. 43–4, 04/01 2012.

[29] A. E. Teschendorff, J. West, and S. Beck, “Age-associated epigenetic drift: implications, and a case of epigenetic thrift?,” Human Molecular Genetics, vol. 22, pp. R7–R15, October 15, 2013 2013.

[30] T. Zhu, S. C. Zheng, D. S. Paul, S. Horvath, and A. E. Teschendorff, “Cell and tissue type independent age-associated DNA methylation changes are not rare but common,” Aging, vol. 10, pp. 3541–3557, 2018.

[31] Y. Guo, Y. Fan, J. Zhang, G. A. Lomberk, Z. Zhou, L. Sun, et al., “Perhexiline activates KLF14 and reduces atherosclerosis by modulating ApoA-I production,” J Clin Invest, vol. 125, pp. 3819–30, Oct 1 2015.

[32] C. Iwaya, H. Kitajima, K. Yamamoto, Y. Maeda, N. Sonoda, H. Shibata, et al., “DNA methylation of the Klf14 gene region in whole blood cells provides prediction for the chronic inflammation in the adipose tissue,” Biochemical and biophysical research communications, vol. 497, pp. 908–915, 2018/03// 2018.

[33] S. E. Jung, S. M. Lim, S. R. Hong, E. H. Lee, K. J. Shin, and H. Y. Lee, “DNA methylation of the ELOVL2, FHL2, KLF14, C1orf132/MIR29B2C, and TRIM59 genes for age prediction from blood, saliva, and buccal swab samples,” Forensic Sci Int Genet, vol. 38, pp. 1–8, Jan 2019.

[34] L. Kananen, S. Marttila, T. Nevalainen, J. Jylhävä, N. Mononen, M. Kähönen, et al., “Aging-associated DNA methylation changes in middle-aged individuals: the Young Finns study,” BMC Genomics, vol. 17, p. 103, Feb 9 2016.

[35] Y. Zhang, X. Yang, B. Gui, G. Xie, D. Zhang, Y. Shang, et al., “Corepressor protein CDYL functions as a molecular bridge between polycomb repressor complex 2 and repressive chromatin mark trimethylated histone lysine 27,” J Biol Chem, vol. 286, pp. 42414–25, Dec 9 2011.

[36] Y. Liu, S. Liu, S. Yuan, H. Yu, Y. Zhang, X. Yang, et al., “Chromodomain protein CDYL is required for transmission/restoration of repressive histone marks,” J Mol Cell Biol, vol. 9, pp. 178–194, Jun 1 2017.

[37] T. R. Robeck, Z. Fei, A. T. Lu, A. Haghani, E. Jourdain, J. A. Zoller, et al., “Multi-species and multi-tissue methylation clocks for age estimation in toothed whales and dolphins,” Commun Biol, vol. 4, p. 642, May 31 2021.

[38] J. A. Anderson, R. A. Johnston, A. J. Lea, F. A. Campos, T. N. Voyles, M. Y. Akinyi, et al., “High social status males experience accelerated epigenetic aging in wild baboons,” eLife, vol. 10, p. e66128, 2021.

[39] N. E. Schlabritz-Loutsevitch, C. J. Dudley, J. J. Gomez, C. H. Nevill, B. K. Smith, S. L. Jenkins, et al., “Metabolic adjustments to moderate maternal nutrient restriction,” British journal of nutrition, vol. 98, pp. 276–284, 2007.

[40] J. V. Kavitha, F. J. Rosario, M. J. Nijland, T. J. McDonald, G. Wu, Y. Kanai, et al., “Down-regulation of placental mTOR, insulin/IGF-I signaling, and nutrient transporters in response to maternal nutrient restriction in the baboon,” FASEB journal: official publication of the Federation of American Societies for Experimental Biology, vol. 28, pp. 1294–1305, 2014.

[41] N. E. Schlabritz-Loutsevitch, K. Howell, K. Rice, E. J. Glover, C. H. Nevill, S. L. Jenkins, et al., “Development of a system for individual feeding of baboons maintained in an outdoor group social environment,” Journal of Medical Primatology, vol. 33, pp. 117–126, 2004/06/01 2004.

[42] S. M. Zehr, R. G. Roach, D. Haring, J. Taylor, F. H. Cameron, and A. D. Yoder, “Life history profiles for 27 strepsirrhine primate taxa generated using captive data from the Duke Lemur Center,” Scientific Data, vol. 1, p. 140019, 2014/07/22 2014.

[43] S. Morgello, B. Gelman, P. Kozlowski, H. Vinters, E. Masliah, M. Cornford, et al., “The National NeuroAIDS Tissue Consortium: a new paradigm in brain banking with an emphasis on infectious disease.,” Neuropathol Appl Neurobiol, vol. 27, pp. 326–35., 2001.

[44] S. Horvath, D. J. Stein, N. Phillips, S. J. Heany, M. S. Kobor, D. T. S. Lin, et al., “Perinatally acquired HIV infection accelerates epigenetic aging in South African adolescents,” AIDS (London, England), vol. 32, pp. 1465–1474, 2018.

[45] S. Horvath and B. R. Ritz, “Increased epigenetic age and granulocyte counts in the blood of Parkinson’s disease patients,” Aging (Albany NY), vol. 7, pp. 1130–42, Dec 2015.

[46] S. Kabacik, S. Horvath, H. Cohen, and K. Raj, “Epigenetic ageing is distinct from senescence-mediated ageing and is not prevented by telomerase expression,” Aging (Albany NY), vol. 10, pp. 2800–2815, Oct 17 2018.

[47] W. Zhou, T. J. Triche, Jr, P. W. Laird, and H. Shen, “SeSAMe: reducing artifactual detection of DNA methylation by Infinium BeadChips in genomic deletions,” Nucleic Acids Research, vol. 46, pp. e123–e123, 2018.

[48] J. Friedman, T. Hastie, and R. Tibshirani, “Regularization Paths for Generalized Linear Models via Coordinate Descent,” Journal of Statistical Software, vol. 33, pp. 1–22, 2010.

[49] T. R. Dawber, G. F. Meadors, and F. E. Moore, Jr., “Epidemiological approaches to heart disease: the Framingham Study,” Am J Public Health Nations Health, vol. 41, pp. 279–81, Mar 1951.

[50] S. Horvath, M. Gurven, M. E. Levine, B. C. Trumble, H. Kaplan, H. Allayee, et al., “An epigenetic clock analysis of race/ethnicity, sex, and coronary heart disease,” Genome Biol, vol. 17, p. 171, 2016.

[51] D. A. Bennett, J. A. Schneider, Z. Arvanitakis, and R. S. Wilson, “Overview and findings from the religious orders study,” Curr Alzheimer Res, vol. 9, pp. 628–45, Jul 2012.

[52] C. Y. McLean, D. Bristor, M. Hiller, S. L. Clarke, B. T. Schaar, C. B. Lowe, et al., “GREAT improves functional interpretation of cis-regulatory regions,” Nat Biotechnol, vol. 28, 2010// 2010.

