## Supplementary Figures and Tables for "Pan-primate DNA methylation clocks"

### SUPPLEMENTARY MATERIAL

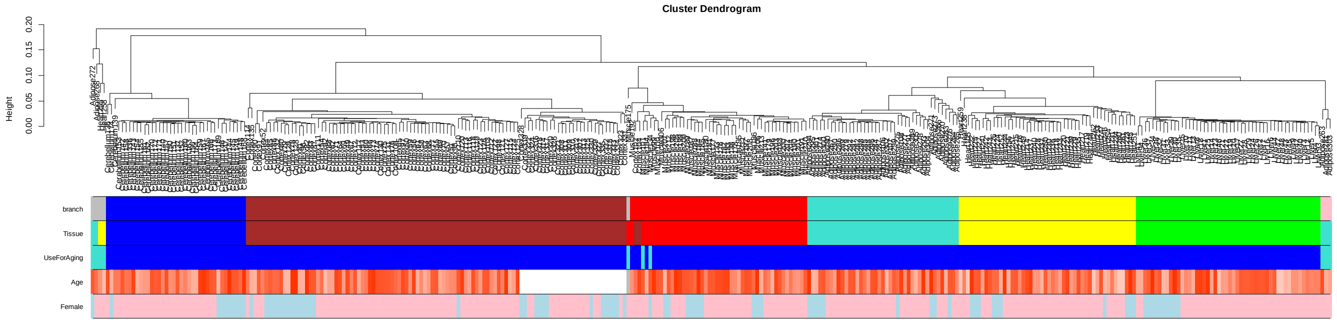


**Supplementary Figure S1**. Unsupervised hierarchical clustering of tissue samples from baboons. Average linkage hierarchical clustering based on the interarray correlation coefficient (Pearson correlation). A height cut-off of 0.07 led to branch colors that largely correspond to Tissue type (second panel): cerebral cortex=brown, muscle=red, heart=yellow, liver=green, adipose=turquoise, cerebellum=blue. A handful of outlying arrays were removed from the analysis (turquoise color in the third color band). Branches largely correspond to tissue type as one can see by comparing the first two color bands.

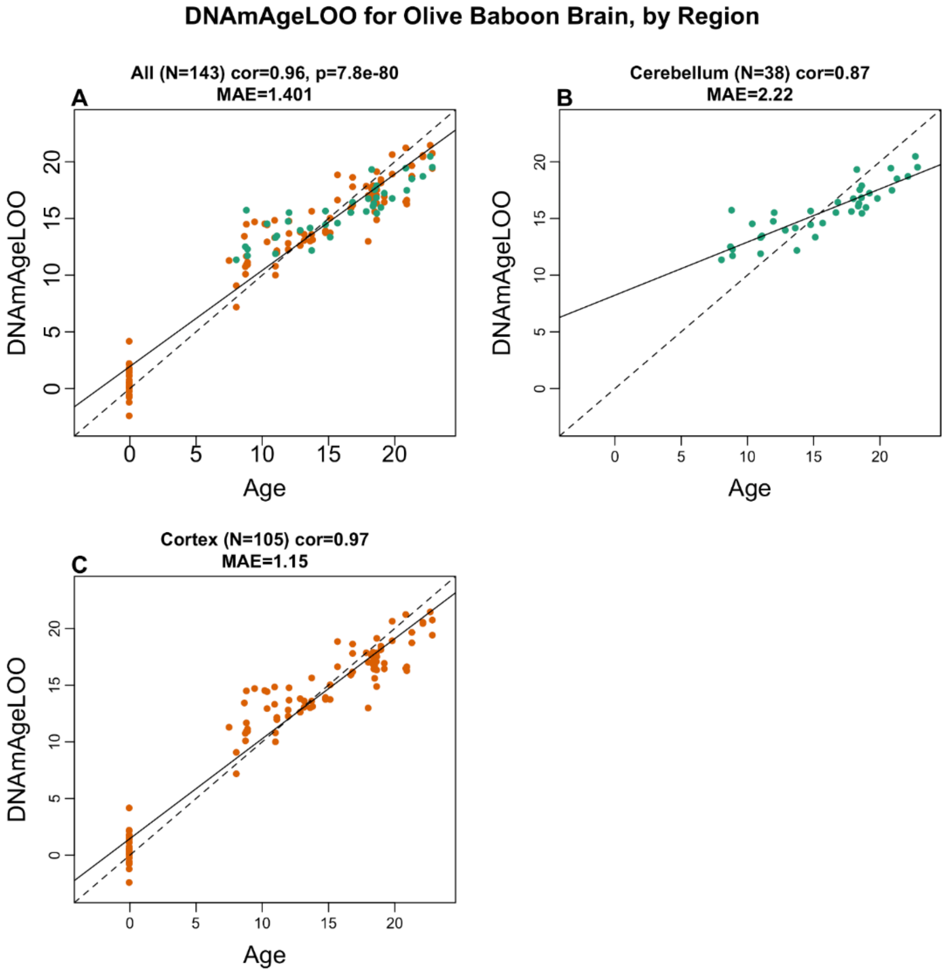


**Supplementary Figure S2. Baboon brain clock** Leave-one-sample-out estimate of age based on DNA methylation data (y-axis) versus chronological age (in units of years). A) The brain clock was developed using cerebellum, frontal cortex and temporal cortex samples. Results restricted to B) cerebellar samples and C) cortical samples Each title reports the sample size, Pearson correlation coefficient and median absolute deviation (median error).

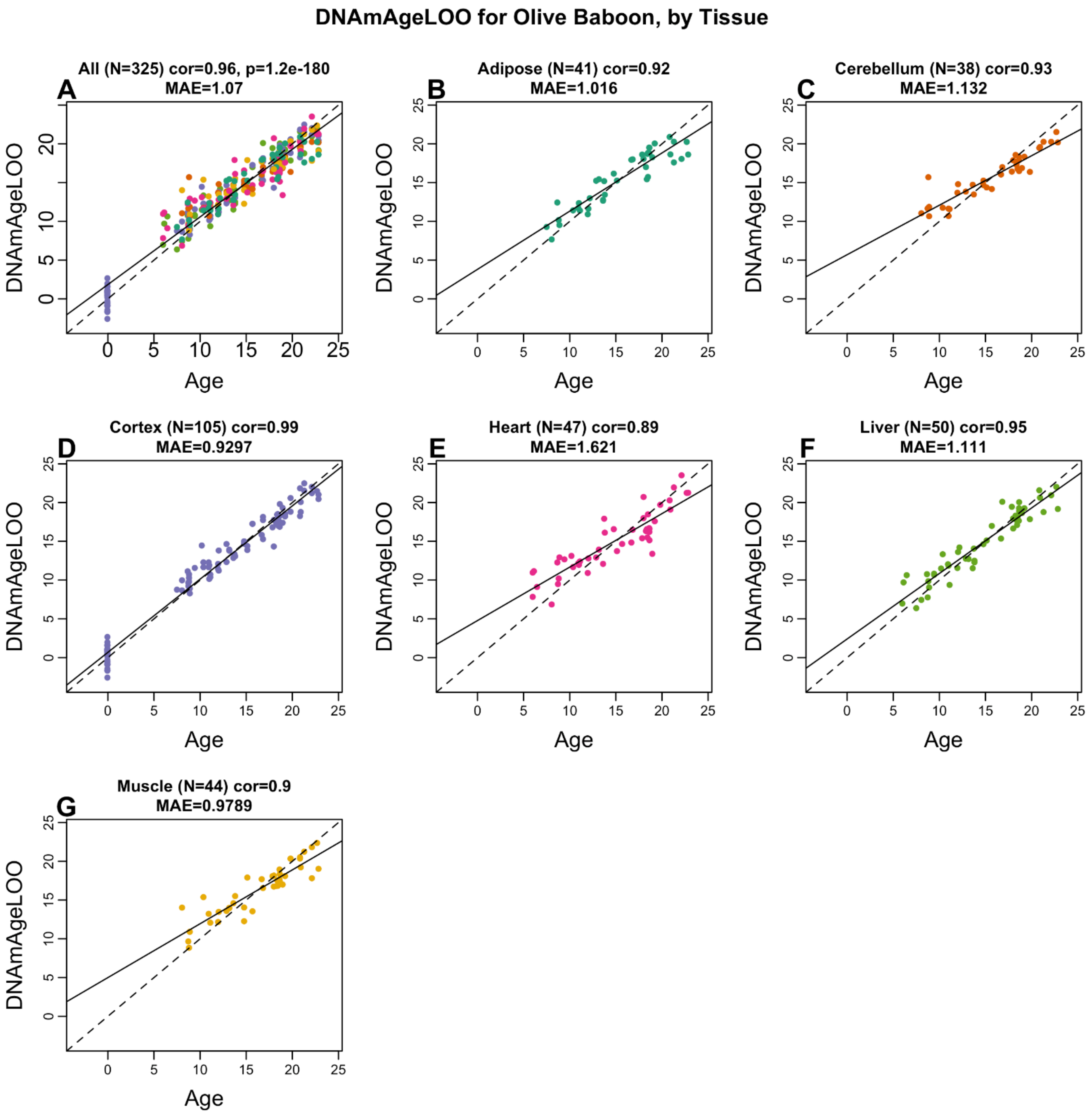


**Supplementary Figure S3. Pan-tissue baboon epigenetic clock**. Leave-one-sample-out estimate of age based on DNA methylation data (y-axis) versus chronological age (in units of years). Results for A) all tissues, B) adipose, C) brain cerebellum, D) cerebellar cortex, E) heart, F) liver, G) skeletal muscle. Each panel reports the sample size, Pearson correlation coefficient and median absolute deviation (median error).

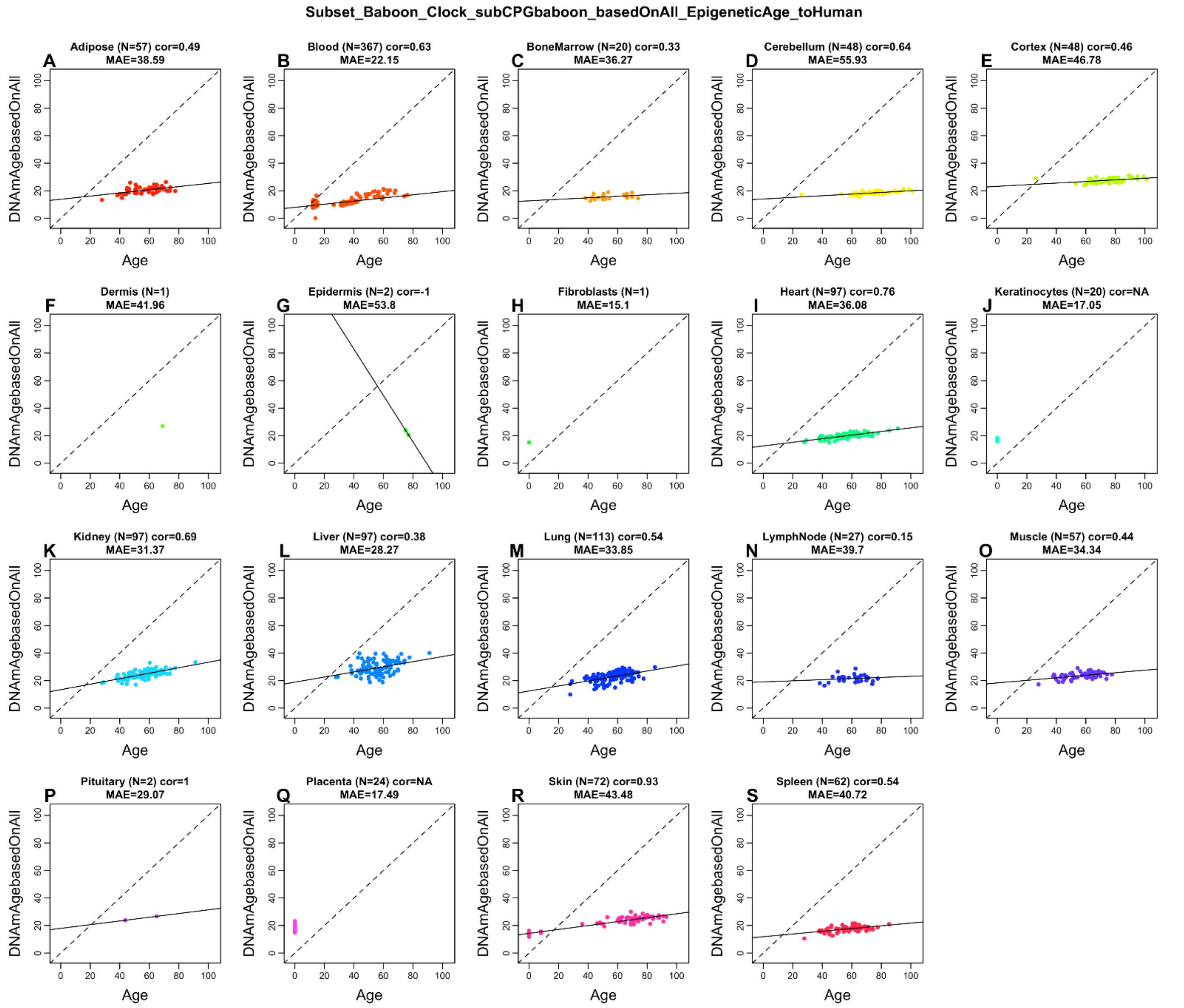


**Supplementary Figure S4. Pan-tissue clock from baboons applied to human tissues**. Baboon pan-tissue clock (trained on baboon tissues) applied to human A) adipose, B) blood, C) bone marrow, D) dermis, E) epidermis, F) fibroblasts G) heart, H) keratinocytes, I) kidney, J) liver, K) lung, L) lymph node, M) muscle, N) pituitary gland, O) skin, P) spleen. Estimate age based on the baboon pan tissue clock (y-axis) versus chronological age in humans (in units of years). Title: sample size, Pearson correlation coefficient and median absolute deviation (median error).

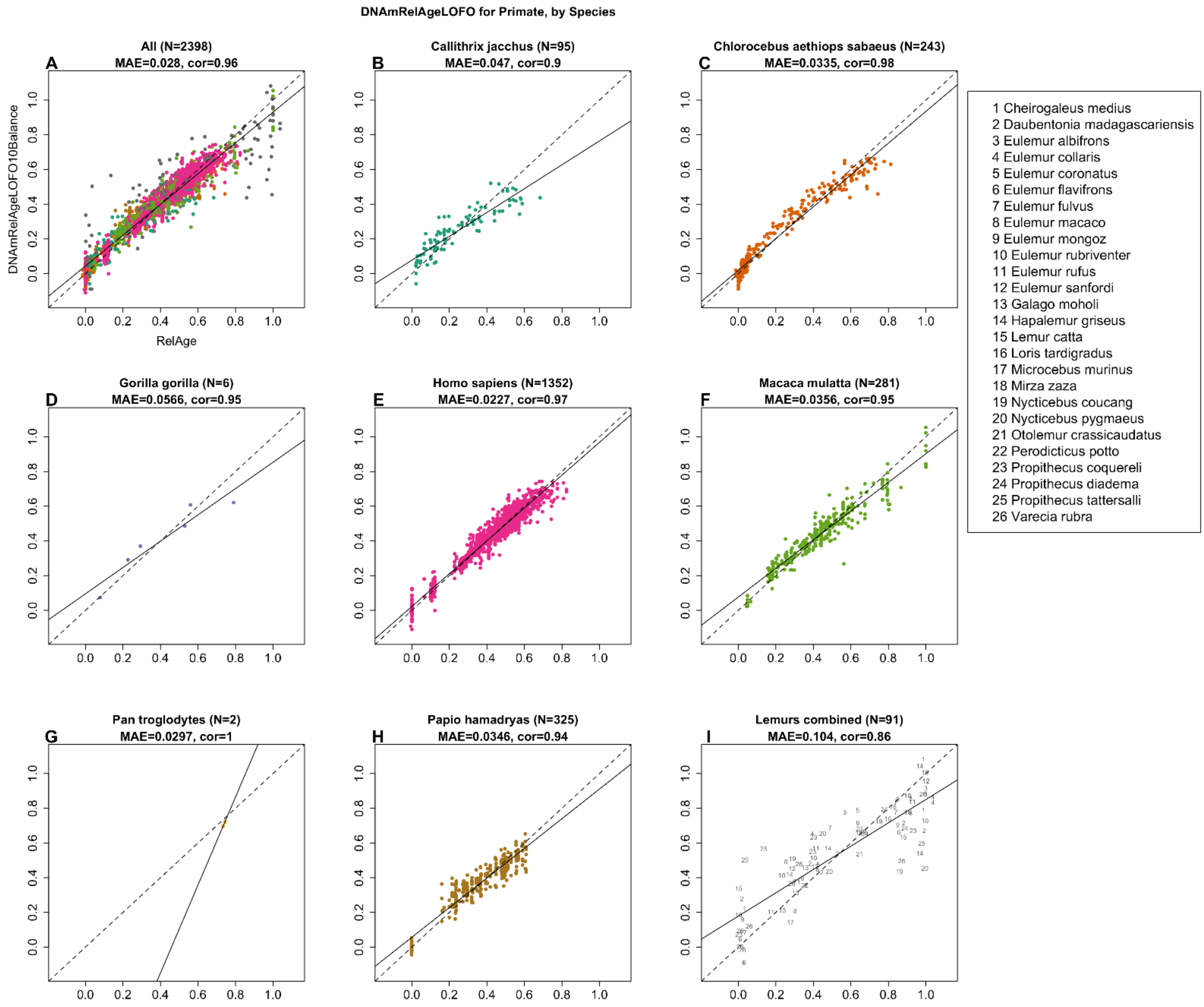


**Supplementary Figure S5. Primate clock for relative age.** A) Epigenetic clock based on tissues from all primate species (colored by species as indicated in the other panels). B-I) are excerpts from panel A but restricted to specific species mentioned in the title. I) Samples (dots) are blood and skin samples from 26 species of strepsirrhines as indicated in legend.
Relative age is defined as the ratio of chronological age with the maximum lifespan of the species.
The maximum lifespan of each species is reported in a Supplementary Table. Relative age (x-axis) versus the ten-fold cross validation (balanced by species) of relative age based on methylation (y-axis). Each panel reports the sample size, correlation coefficient, median absolute error (MAE). Dots are colored by species. The primate clock was developed by regressing relative age on cytosines that map to baboons and humans. Details can be found in the Supplement.

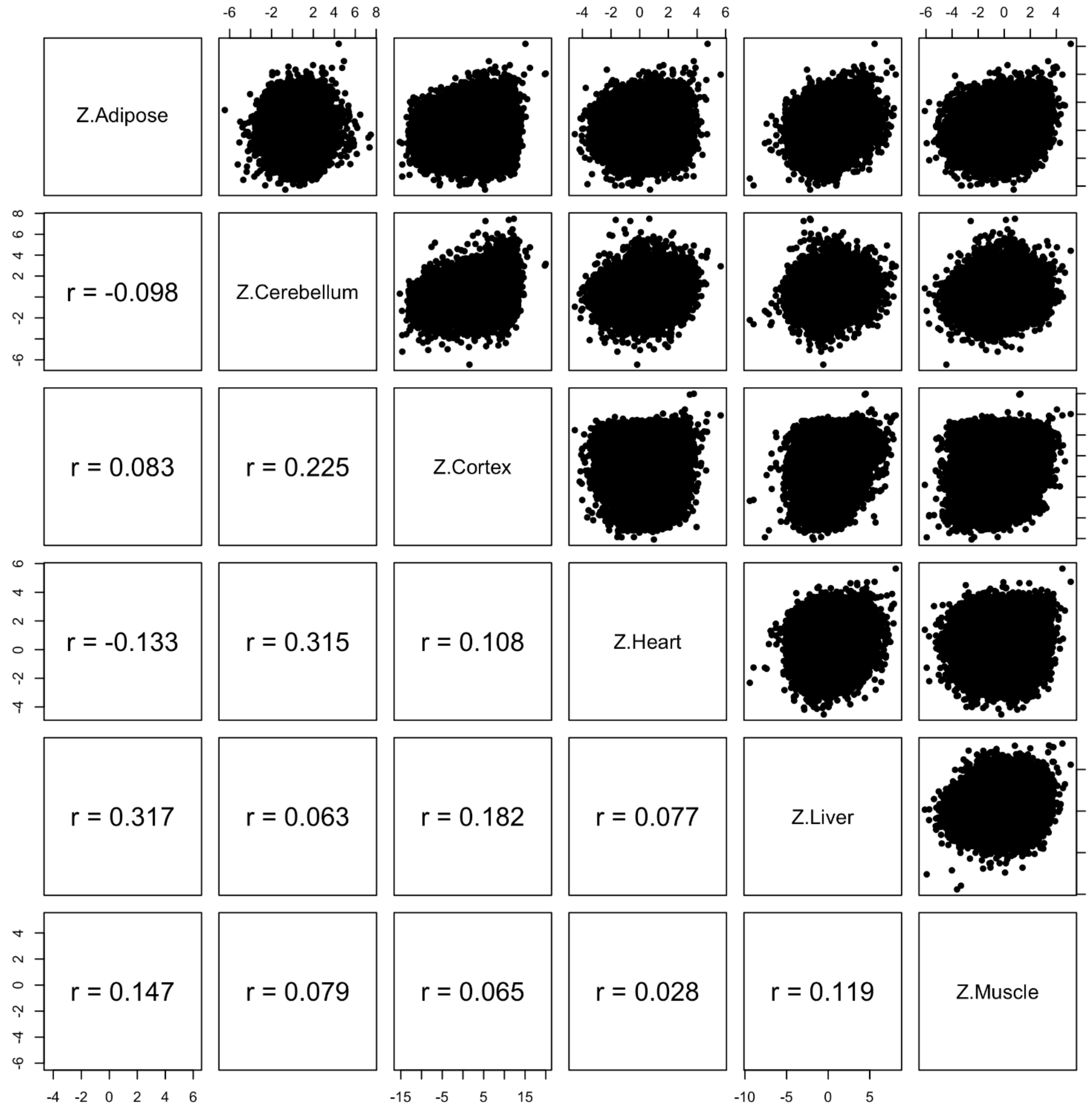


**Supplementary Figure S6. Epigenome wide association study of correlation in different baboon tissues**. Each dot corresponds to a CpG. Z statistics for a correlation test of age in adipose, cerebellum, brain cortex, heart, liver, muscle, temporal cortex.

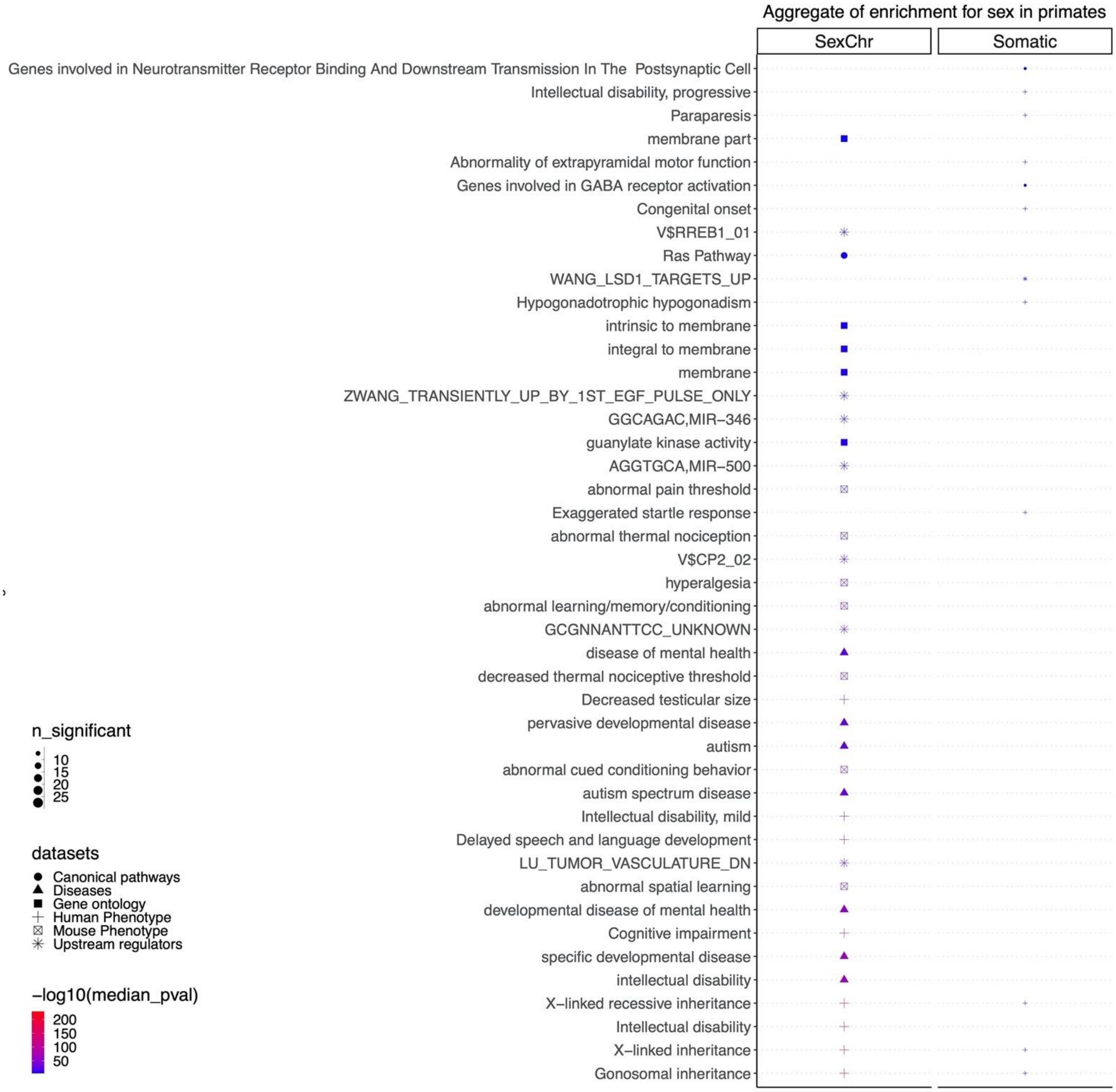


**Supplementary Figure S7. Enrichment analysis of sex-related CpGs in primate tissues by sex chromosomes**. The figure summarizes the results of 27 EWAS of sex corresponding to 27 different strata comprised by primate species and tissue type. For each term (y-axis) we obtained 27 enrichment p values (hypergeometric test) corresponding to the 27 strata. These 27 p values were summarized by their median value. This unusual meta analysis approach leads to a *descriptive* measure of significance as opposed to an inferential measure. The size of the symbol (n_significant) corresponds to the number of times that a term was significant ($p<0.001$) across the 27 analyzed datasets. The top three enriched datasets from each category (Canonical pathways, diseases, gene ontology, human and mouse phenotypes, and upstream regulators) were selected and further filtered for significance at median pvalue $< 10^{-5}$. We excluded marmosets from the analysis for reasons mentioned in the text.

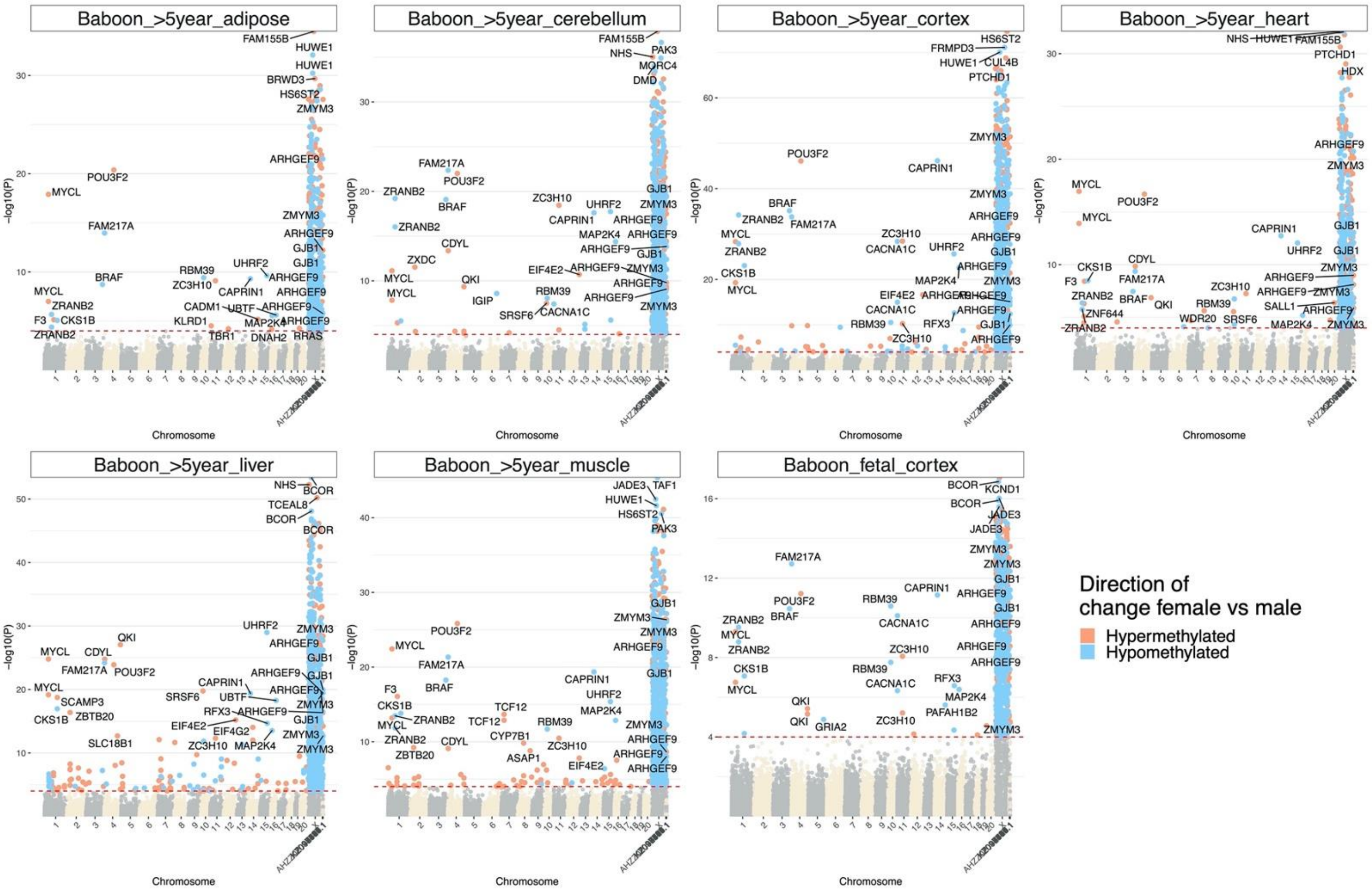

**Supplementary Figure S8. Manhattan plots of sex difference in baboon samples**. The coordinates are estimated based on the alignment of Mammalian array probes to Panu_3.0.100 genome assembly. The direction of change for female vs male with $p < 10^{-4}$ (red dotted line) is highlighted by red (increased methylation in females compared to males) and blue (decreased methylation in females) colors. The top 5 X chromosomal CpGs, and top 30 non-X chromosomal markers (i.e. autosomal) CpGs were labeled by neighboring genes.

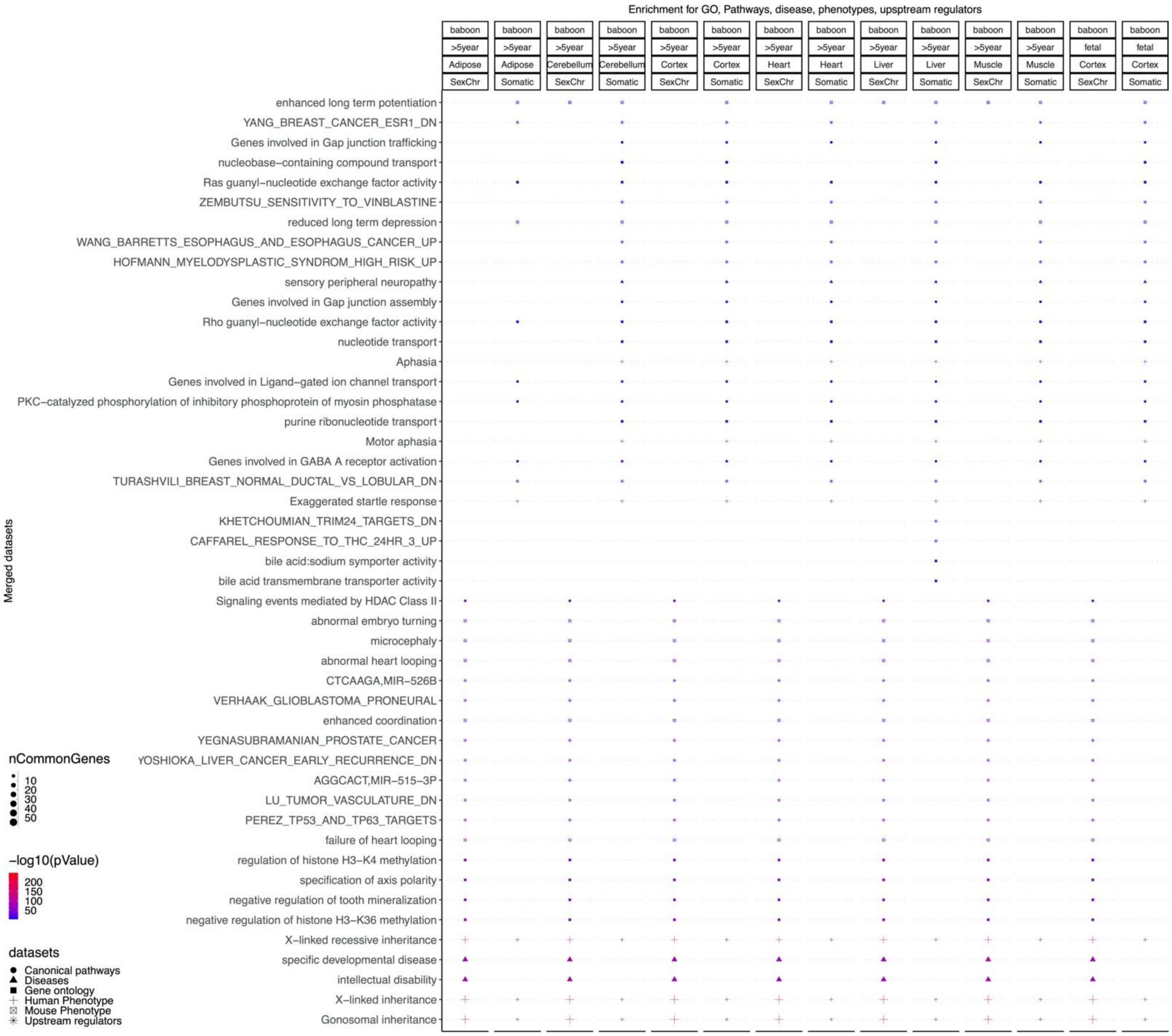


**Supplementary Figure S9. Enrichment analysis of the top CpGs associated with sex in baboons.** The gene level enrichment was done using GREAT analysis and human Hg19 background. The background probes were limited to probes that were mapped to the same gene in the olive baboon genome. The top two enriched datasets from each category (Canonical pathways, diseases, gene ontology, human and mouse phenotypes, and upstream regulators) were selected and further filtered for significance at $p < 10^{-10}$.

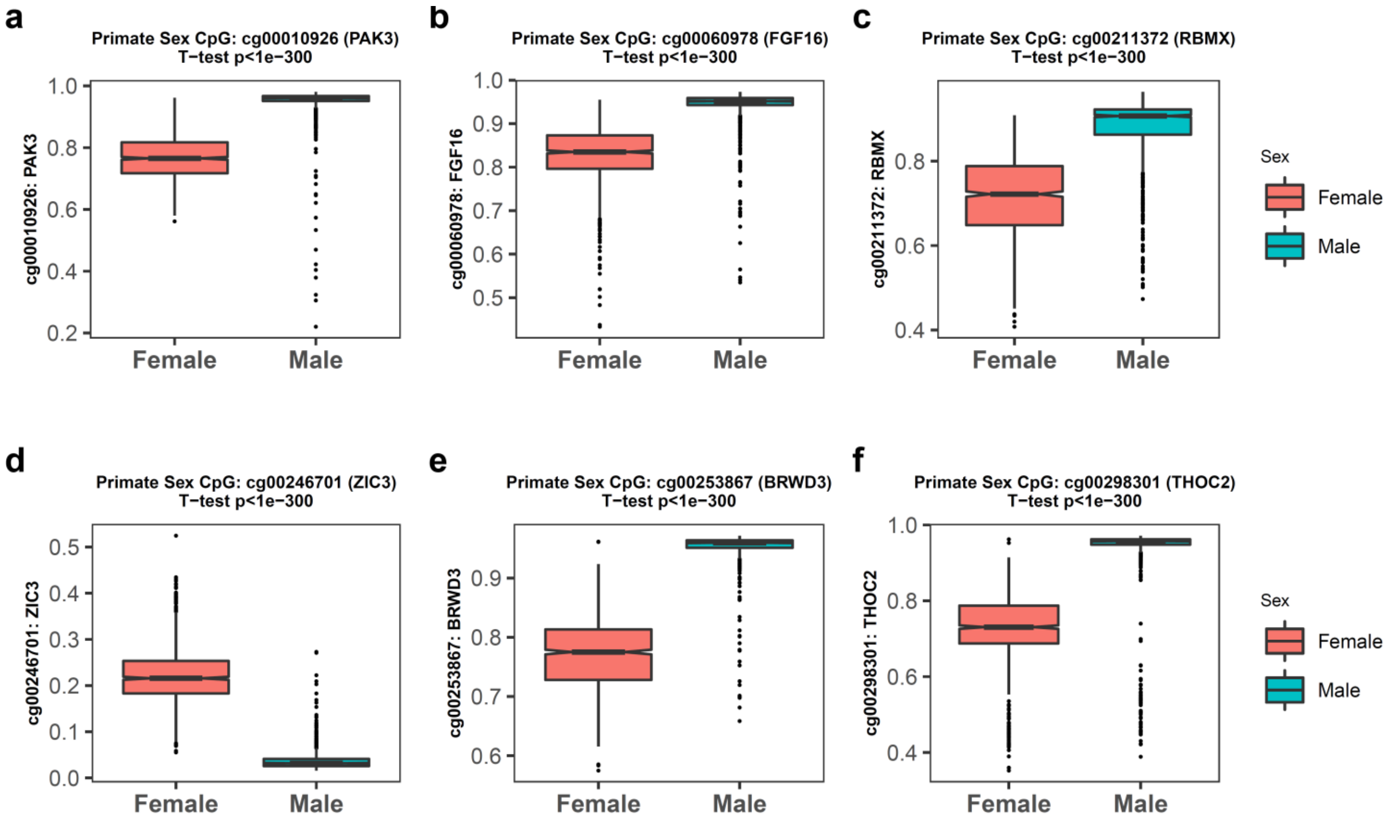


**Supplementary Figure S10.** Select CpGs that are highly associated with sex in primates. Each panel presents a CpG (and adjacent gene) that is highly associated with sex across all primate species. Student T-test p values.

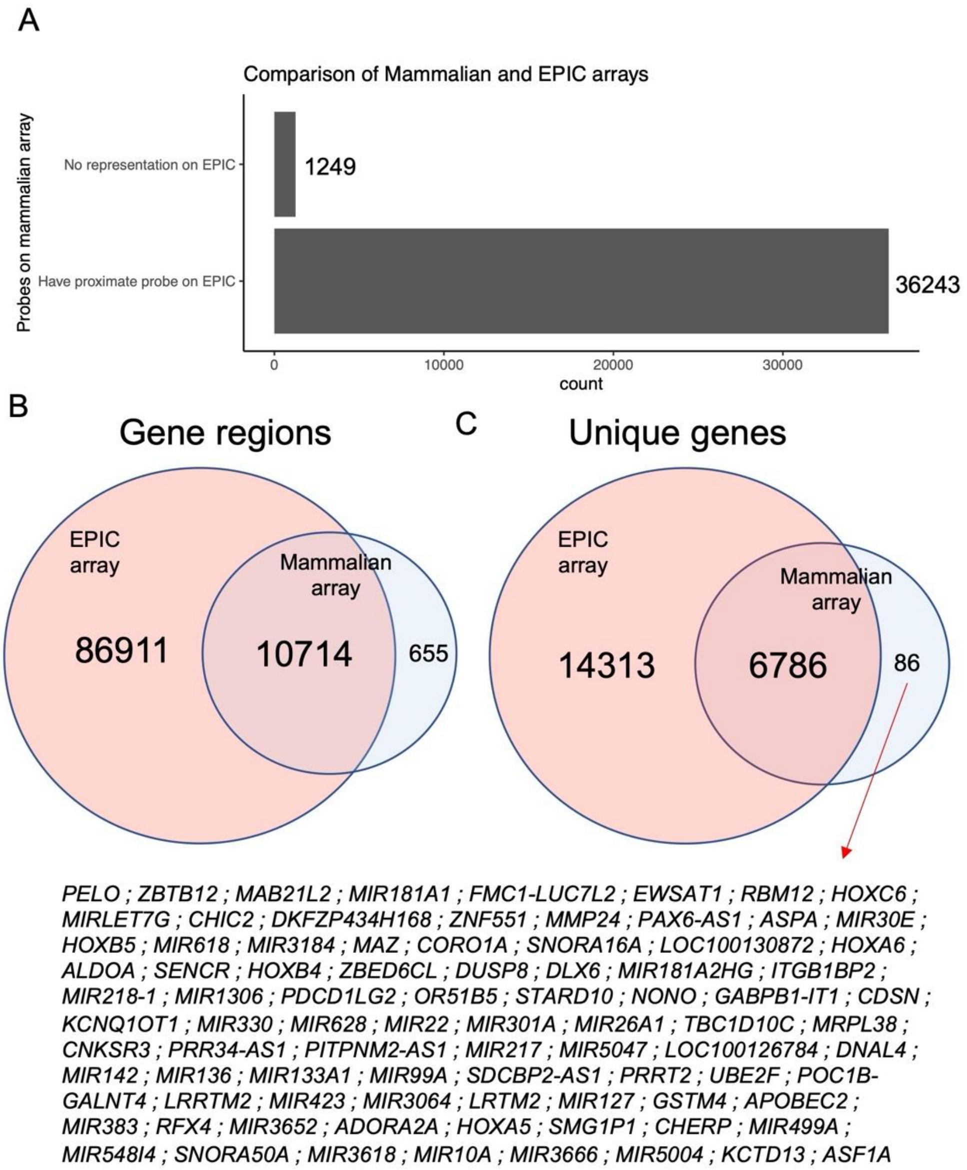


**Supplementary Figure S11. Mammalian array compared to the human Illumina EPIC array.** The EPIC array covers most of the genomic regions represented on mammalian array. A) Most of the mammalian array probes are located on gene regions that have at least one probe represented on the EPIC array. B) Venn Diagram visualizing the overlap between gene regions covered by the EPIC array and the mammalian array. Most of the gene regions on the mammalian array are also covered in EPIC array. C) Venn Diagram visualizing the overlap between genes covered by the EPIC array and the mammalian array. There are 86 genes that are specifically presented on mammalian but not the EPIC array.

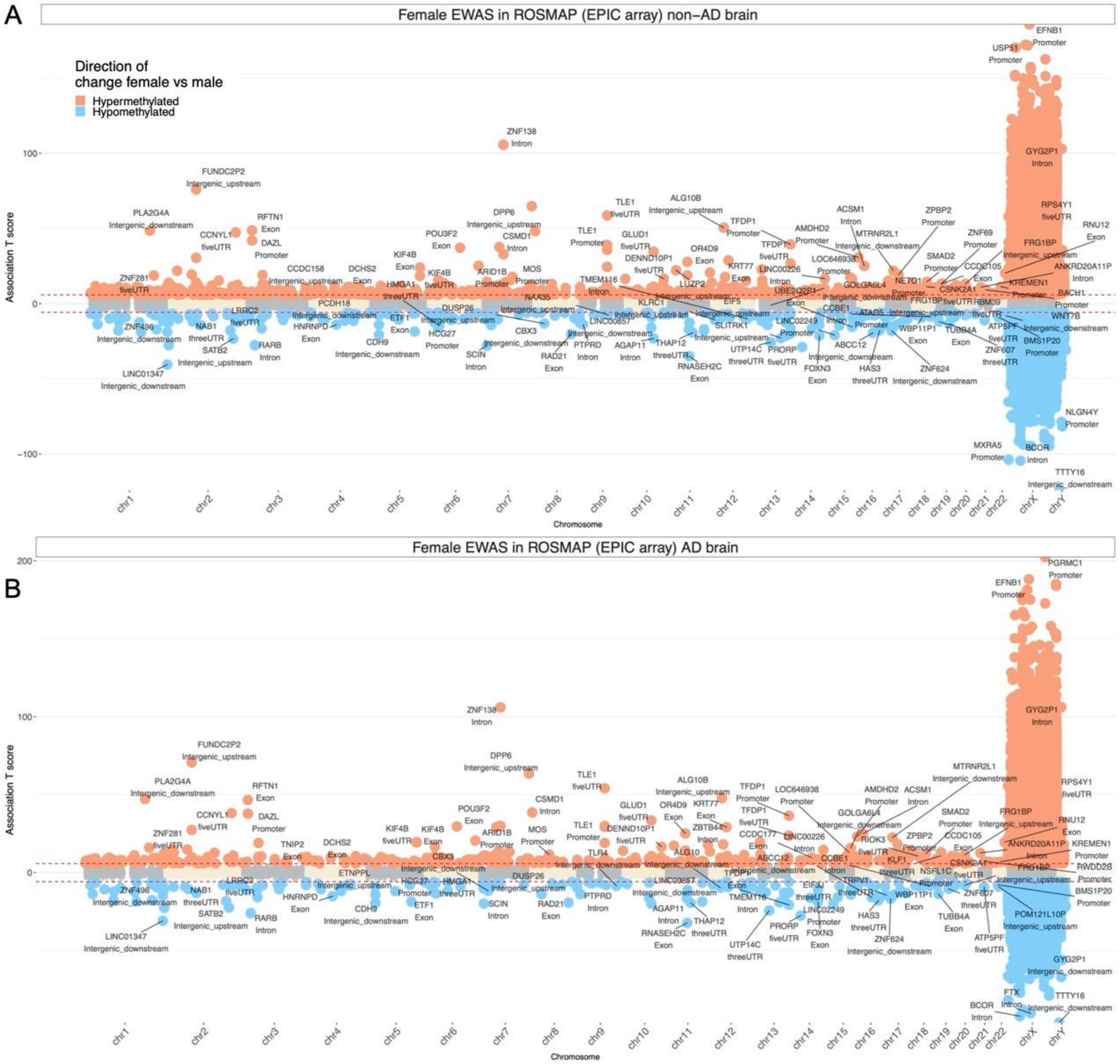


**Supplementary Figure S12. Sex differences in DNAm pattern in human postmortem prefrontal cortex samples.** Results for individuals with (A) and without (B) Alzheimer's disease (AD) neuropathology. Sex differences were adjusted for age in the model. The red lines represent T scores at $p < 10^{-8}$.
The EWAS results of sex are based on human, baboon, Baboon, rhesus macaque.

### Supplementary Tables

| Row | Species | CommonName | N | N.Female | N.Blood | N.Skin | N.Liver | N.Cortex | Age.Min | Age.Med | Age.Max |
|---|---|---|---|---|---|---|---|---|---|---|---|
| 1 | Callithrix geoffroyi | White-fronted marmoset | 1 | 1 | 1 | 0 | 0 | 0 | 18.2 | 18.2 | 18.2 |
| 2 | Callithrix jacchus | Common marmoset | 95 | 45 | 95 | 0 | 0 | 0 | 0.5 | 4.4 | 15.6 |
| 3 | Cheirogaleus medius | Fat-tailed dwarf lemur | 3 | 0 | 2 | 1 | 0 | 0 | 1.0 | 29.6 | 29.6 |
| 4 | Chlorocebus aethiops sabaeus | Vervet monkey | 243 | 152 | 144 | 0 | 48 | 48 | -0.4 | 3.1 | 25.0 |
| 5 | Daubentonia madagascariensis | Aye-aye | 4 | 3 | 3 | 1 | 0 | 0 | 0.7 | 23.4 | 36.7 |
| 6 | Eulemur albifrons | White-headed lemur | 3 | 2 | 2 | 1 | 0 | 0 | 19.7 | 34.6 | 34.6 |
| 7 | Eulemur collaris | Collared brown lemur | 4 | 3 | 3 | 1 | 0 | 0 | 0.8 | 23.4 | 33.8 |
| 8 | Eulemur coronatus | Crowned lemur | 3 | 0 | 2 | 1 | 0 | 0 | 1.0 | 12.8 | 19.1 |
| 9 | Eulemur flavifrons | Blue-eyed black lemur | 4 | 0 | 3 | 1 | 0 | 0 | 0.3 | 17.0 | 27.4 |
| 10 | Eulemur fulvus | Brown lemur | 3 | 2 | 2 | 1 | 0 | 0 | 17.4 | 29.8 | 29.8 |
| 11 | Eulemur macaco | Black lemur | 4 | 1 | 3 | 1 | 0 | 0 | 9.5 | 22.9 | 34.5 |
| 12 | Eulemur mongoz | Mongoose lemur | 4 | 2 | 3 | 1 | 0 | 0 | 0.8 | 21.6 | 30.8 |
| 13 | Eulemur rubriventer | Red-bellied lemur | 4 | 3 | 3 | 1 | 0 | 0 | 7.9 | 23.7 | 33.8 |
| 14 | Eulemur rufus | Red lemur | 3 | 1 | 3 | 0 | 0 | 0 | 5.7 | 13.5 | 30.3 |
| 15 | Eulemur sanfordi | Sanford's brown lemur | 4 | 3 | 3 | 1 | 0 | 0 | 9.4 | 23.6 | 32.9 |
| 16 | Galago moholi | South African galago | 3 | 1 | 2 | 1 | 0 | 0 | 3.9 | 5.9 | 6.8 |
| 17 | Gorilla gorilla | Gorilla | 3 | 1 | 3 | 0 | 0 | 0 | 4.7 | 13.6 | 17.6 |
| 18 | Gorilla gorilla | Western lowland gorilla | 3 | 1 | 3 | 0 | 0 | 0 | 31.8 | 33.6 | 47.5 |
| 19 | Hapalemur griseus | Bamboo lemur | 4 | 3 | 3 | 1 | 0 | 0 | 7.4 | 19.5 | 26.1 |
| 20 | Homo sapiens | Human | 1352 | 655 | 511 | 74 | 97 | 46 | -0.8 | 53.7 | 101.0 |
| 21 | Lemur catta | Ring-tailed lemur | 4 | 3 | 3 | 1 | 0 | 0 | 0.1 | 5.7 | 32.8 |

| | | | | | | | | | | | |
|---|---|---|---|---|---|---|---|---|---|---|---|
| 22 | Loris tardigradus | Slender loris | 2 | 1 | 1 | 1 | 0 | 0 | 17.2 | 17.5 | 17.8 |
| 23 | Macaca mulatta | Rhesus macaque | 281 | 99 | 199 | 51 | 5 | 6 | 1.8 | 18.5 | 42.0 |
| 24 | Microcebus murinus | Gray mouse lemur | 2 | 1 | 2 | 0 | 0 | 0 | 0.5 | 2.8 | 5.1 |
| 25 | Mirza zaza | Northern giant mouse lemur | 3 | 0 | 2 | 1 | 0 | 0 | 12.9 | 18.1 | 18.1 |
| 26 | Nycticebus coucang | Slow loris | 3 | 2 | 2 | 1 | 0 | 0 | 7.4 | 19.3 | 22.2 |
| 27 | Nycticebus pygmaeus | Pygmy slow loris | 4 | 2 | 3 | 1 | 0 | 0 | 0.7 | 9.3 | 19.9 |
| 28 | Otolemur crassicaudatus | Greater galago | 4 | 4 | 3 | 1 | 0 | 0 | 7.1 | 13.4 | 14.7 |
| 29 | Pan troglodytes | Chimpanzee | 2 | 2 | 2 | 0 | 0 | 0 | 43.6 | 43.9 | 44.2 |
| 30 | Papio hamadryas | Olive baboon | 325 | 248 | 0 | 0 | 50 | 105 | -0.1 | 14.8 | 22.8 |
| 31 | Perodicticus potto | Potto | 1 | 0 | 1 | 0 | 0 | 0 | 11.5 | 11.5 | 11.5 |
| 32 | Pongo pygmaeus | Orangutan | 1 | 1 | 1 | 0 | 0 | 0 | 43.7 | 43.7 | 43.7 |
| 33 | Propithecus coquereli | Potto | 4 | 1 | 3 | 1 | 0 | 0 | 0.1 | 12.2 | 28.5 |
| 34 | Propithecus diadema | Diademed sifaka | 3 | 0 | 2 | 1 | 0 | 0 | 14.2 | 16.4 | 18.7 |
| 35 | Propithecus tattersalli | Golden-crowned sifaka | 3 | 0 | 3 | 0 | 0 | 0 | 3.5 | 17.3 | 25.4 |
| 36 | Varecia rubra | Red ruffed lemur | 8 | 1 | 6 | 2 | 0 | 0 | 0.4 | 6.9 | 39.4 |
| 37 | Varecia variegata | Variegated lemur | 1 | 0 | 1 | 0 | 0 | 0 | 12.0 | 12.0 | 12.0 |

**Supplementary Table S1. Data used for the primate clocks.** Columns report the species (Latin Name, Common Name). Total sample size (N), i.e. the number of arrays/DNA samples per species. Number of samples from females. Number of samples from different tissues. Age: minimum, maximum, median.

| Tissue | N | No. Female | Mean Age | Min. Age | Max. Age |
|---|---|---|---|---|---|
| Adipose | 41 | 32 | 15.6 | 7.49 | 22.8 |
| Cerebellum | 38 | 29 | 15.8 | 8.04 | 22.8 |
| Cortex | 76 | 60 | 15.3 | 7.49 | 22.8 |

| | | | | | |
|---|---|---|---|---|---|
| Fetal Cortex | 29 | 16 | 0 | 0 | 0 |
| Heart | 48 | 38 | 14.5 | 5.98 | 22.8 |
| Liver | 50 | 40 | 14.6 | 5.98 | 22.8 |
| Muscle | 44 | 34 | 16.4 | 8.04 | 22.8 |

**Table S2. Baboon samples.**
Tissue type. N=Total number of samples per tissue. Number of females. Age: mean, minimum and maximum. The fetal brain cortex samples were collected at gestational age 165 days.

| **Latin Name** | **Common Name** | **Avg. Maturity** | **GestationTimeInYeares** | **max. Lifespan (Years)** |
|---|---|---|---|---|
| Callithrix geoffroyi | White-fronted marmoset | 1.375 | 0.407 | 19 |
| Callithrix jacchus | Common marmoset | 1.177 | 0.395 | 22.8 |
| Cheirogaleus medius | Fat-tailed dwarf lemur | 1.000 | 0.167 | 30 |
| Chlorocebus aethiops sabaeus | Vervet | 3.916 | 0.444 | 30.8 |
| Daubentonia madagascariensis | Aye-aye | 2.416 | 0.452 | 37 |
| Eulemur albifrons | White-headed lemur | 8.234 | 0.329 | 34.59 |
| Eulemur collaris | Collared brown lemur | 12.137 | 0.337 | 32.61 |
| Eulemur coronatus | Crowned lemur | 1.666 | 0.345 | 30 |
| Eulemur flavifrons | Blue-eyed black lemur | 2.100 | 0.348 | 32 |
| Eulemur fulvus | Brown lemur | 1.564 | 0.323 | 35.5 |
| Eulemur macaco | Black lemur | 1.104 | 0.329 | 37.5 |
| Eulemur mongoz | Mongoose lemur | 2.349 | 0.329 | 36.2 |
| Eulemur rubriventer | Red-bellied lemur | 2.734 | 0.329 | 34 |
| Eulemur rufus | Red lemur | 12.426 | 0.329 | 32.67 |
| Eulemur sanfordi | Sanford's brown lemur | 7.836 | 0.329 | 32.94 |
| Galago moholi | South African galago | 0.710 | 0.340 | 16.6 |
| Gorilla gorilla | Western lowland gorilla | 9.375 | 0.701 | 60.1 |
| Gorilla gorilla | Gorilla | 9.375 | 0.701 | 60.1 |
| Hapalemur griseus | Bamboo lemur | 3.049 | 0.397 | 27 |
| Homo sapiens | Human | 13.500 | 0.767 | 122.5 |
| Lemur catta | Ring-tailed lemur | 2.064 | 0.370 | 37.3 |
| Loris tardigradus | Slender loris | 1.021 | 0.455 | 21.6 |
| Macaca mulatta | Rhesus macaque | 4.436 | 0.452 | 42 |
| Microcebus murinus | Gray mouse lemur | 0.666 | 0.167 | 18.2 |
| Mirza zaza | Northern giant mouse lemur | 1.100 | 0.245 | 20 |
| Nycticebus coucang | Slow loris | 1.584 | 0.515 | 25.8 |
| Nycticebus pygmaeus | Pygmy slow loris | 0.748 | 0.515 | 20 |
| Otolemur crassicaudatus | Greater galago | 1.553 | 0.356 | 22.7 |
| Pan troglodytes | Chimpanzee | 8.625 | 0.627 | 59.4 |
| Papio hamadryas | Olive baboon | 4.488 | 0.468 | 37.5 |

| | | | | |
|---|---|---|---|---|
| Perodicticus potto | Potto | 1.499 | 0.466 | 32.4 |
| Pongo pygmaeus | Orangutan | 7.000 | 0.682 | 59 |
| Propithecus coquereli | Potto | 3.814 | 0.386 | 30.59 |
| Propithecus diadema | Diademed sifaka | 2.875 | 0.430 | 21 |
| Propithecus tattersalli | Golden-crowned sifaka | 4.499 | 0.404 | 26 |
| Varecia rubra | Red ruffed lemur | 1.725 | 0.268 | 40 |
| Varecia variegata | Red ruffed lemur | 1.718 | 0.268 | 39.4 |

**Supplementary Table S3. Maximum lifespans of 37 primate species used in this article.** Columns report the species common name, species Latin name, primate family, maximum observed lifespan (in years) and average age at sexual maturity (years, averaged across both sexes). These age estimates come from anAge [2] and were updated using the Duke Lemur Center Database - Duke Lemur Center https://lemur.duke.edu/duke-lemur-center-database/ Since its establishment in 1966, the Duke Lemur Center has accumulated detailed records for over 4300 individuals from over 40 closely related yet biologically diverse prosimian primate taxa.

**Supplementary Table S4, S5, S6, S7, S8, S9 can be found in the Excel file.**

### References


[1] C. Y. McLean, D. Bristor, M. Hiller, S. L. Clarke, B. T. Schaar, C. B. Lowe*, et al.*, "GREAT improves functional interpretation of cis-regulatory regions," *Nat Biotechnol,* vol. 28, 2010// 2010.

[2] J. P. de Magalhaes, J. Costa, and G. M. Church, "An analysis of the relationship between metabolism, developmental schedules, and longevity using phylogenetic independent contrasts," *J Gerontol A Biol Sci Med Sci,* vol. 62, pp. 149-60, Feb 2007.

[3] S. Horvath, "DNA methylation age of human tissues and cell types," *Genome Biol,* vol. 14, p. R115, 2013.
